# Single shot dendritic cell targeting SARS-CoV-2 vaccine candidate induces broad and durable systemic and mucosal immune responses

**DOI:** 10.1101/2023.02.21.529344

**Authors:** Nicholas You Zhi Cheang, Peck Szee Tan, Kiren Purushotorma, Wee Chee Yap, Benson Yen Leong Chua, Kai Sen Tan, Kirsteen McInnes Tullett, Aileen Ying-Yan Yeoh, Caris Qi Hui Tan, Xinlei Qian, Douglas Jie Wen Tay, Irina Caminschi, Yee Joo Tan, Paul Anthony Macary, Chee Wah Tan, Mireille Hanna Lahoud, Sylvie Alonso

## Abstract

Current COVID-19 vaccines face certain limitations, which include waning immunity, immune escape by SARS-CoV-2 variants, limited CD8^+^ cellular response, and poor induction of mucosal immunity. Here, we engineered a Clec9A-RBD antibody construct that delivers the Receptor Binding Domain (RBD) from SARS-CoV-2 spike protein to conventional type 1 dendritic cells (cDC1). We showed that single dose immunization with Clec9A-RBD induced high RBD-specific antibody titers with a strong T-helper 1 (T_H_1) isotype profile and exceptional durability, whereby antibody titers were sustained for at least 21 months post-vaccination. Uniquely, affinity maturation of the antibody response was observed over time, as evidenced by enhanced neutralization potency and breadth across the sarbecovirus family. Consistently and remarkably, RBD-specific T-follicular helper cells and germinal center B cells were still detected at 12 months post-immunization. Increased antibody-dependent cell-mediated cytotoxicity (ADCC) activity of the immune sera was also measured over time with comparable efficacy against ancestral SARS-CoV-2 and variants, including Omicron. Furthermore, Clec9A-RBD immunization induced a durable poly-functional T_H_1-biased cellular response that was strongly cross-reactive against SARS-CoV-2 variants, including Omicron, and with robust CD8^+^ T cell signature. Lastly, Clec9A-RBD single dose systemic immunization primed effectively RBD-specific cellular and humoral mucosal immunity in lung. Taken together, Clec9A-RBD immunization has the potential to trigger robust and sustained, systemic and mucosal immune responses against rapidly evolving SARS-CoV2 variants.

## INTRODUCTION

COVID-19 is an acute respiratory viral disease caused by SARS-CoV-2, with over 600 million infections and 6.5 million deaths worldwide as of December 2022 [1, 2]. In severe cases, respiratory failure, pneumonia, and cardiovascular dysfunctions have been observed [3, 4]. Long-term effects of COVID-19 (aka long COVID) affecting respiratory, cardiovascular, digestive, motor, and cognitive activities have also been increasingly recognized as a public health concern [5]. Therefore, safe and effective vaccines are imperative to protect not only against death but also against long term, debilitating conditions associated with SARS-CoV2 infection. Currently, over 13 million doses of COVID-19 vaccines from different platforms have been administered globally [1]. However, they face a number of challenges that limit their effectiveness in conferring potent and long-lasting immunity. Firstly, repeated booster doses are needed as SARS-CoV-2-reactive antibodies and T cells waned over time after vaccination [6, 7]. Secondly, some vaccine platforms induce poor CD8^+^ T cell responses, which have been associated with lower protection against severe disease [6, 8]. Thirdly, immune imprinting has been proposed to limit the ability of vaccines based on full-length spike to protect against SARS-CoV-2 variants with immune escape mutations [9, 10]. Lastly, current vaccines fail to elicit adequate mucosal immunity, which is crucial for protection against re-infection [11]. Thus, new generation vaccines that can overcome at least some of these limitations are needed to provide broad and durable protection against rapidly evolving SARS-CoV-2 variants.

In recent years, targeting antigens to dendritic cells (DC) has emerged as an attractive vaccine strategy that generally improves vaccine effectiveness, and allows antigen and dose sparing [12]. Specifically, targeting antigen vaccine candidates to Clec9A expressed on type I conventional DC (cDC1) subset, has proven promising in pre-clinical studies with strong and sustained humoral and cellular immune responses upon single dose immunization [13-17]. Clec9A is a C-type lectin-like receptor that acts as a damage recognition receptor; Clec9A recognizes filamentous actin exposed when the cell membrane is damaged and facilitates the cross-presentation of dead-cell associated antigens [18, 19]. Clec9A targeting involves an anti-Clec9A monoclonal antibody (mAb) where the vaccine antigen candidate is genetically fused to the C-terminal end of the heavy chains, enabling the targeted delivery of antigens to Clec9A on cDC1. Clec9A targeting was found to be more potent than targeting of other DC receptors [13, 14, 17]. Thanks to high Toll-like receptor 3 (TLR3) expression and potent Type-I interferon (IFN) signaling, cDC1 display enhanced antigen cross-presentation activity, thereby leading to potent CD8^+^ T cell responses to eliminate infected or cancer cells [14, 17]. Moreover, high interleukin-12 (IL-12) secretion by cDC1 drives CD4^+^ T-helper 1 (T_H_1) polarization, promoting the generation of antiviral cytokines, cytotoxic T cell activity, and CD8^+^ memory T cells, which have been correlated with reduced pathophysiology during infection [20, 21]. Furthermore, in view of the restricted expression of Clec9A on cDC1, Clec9A antibody constructs were found to persist in the systemic circulation for a long duration compared to non-targeting and other DC-targeting approaches [13, 15]. This prolonged antigen availability facilitates persistent antigen presentation and T cell stimulation, favoring the production of antigen-specific T-follicular helper (T_FH_) cells, which in turn support germinal centre reactions where B cells undergo affinity maturation [13, 22, 23].

Here, ancestral SARS-CoV-2 Receptor Binding Domain (RBD) was genetically fused to the heavy chains of Clec9A mAb. RBD contains majority of the spike protein’s neutralizing epitopes and has been proposed to limit immune imprinting against SARS-CoV-2 variants [24]. In this study, we characterized the humoral and cellular immune responses triggered in mice upon single dose systemic immunization with Clec9A-RBD antibody construct. Our results indicate that targeting RBD to Clec9A represents an attractive alternative vaccine strategy that induces sustained humoral and cellular, systemic, and mucosal immune responses that react against all SARS-CoV-2 variants.

## RESULTS

### Sustained antibody response and prolonged affinity maturation upon single dose immunization with Clec9A-RBD

We first determined that 2 µg of Clec9A-RBD adjuvanted with poly I:C represented the optimal dose-adjuvant formulation that induced the highest anti-RBD neutralizing titers in adult Balb/c mice upon a single dose **(Figure S1)**. Furthermore, we showed that immunization with an equivalent antigen dose of non-targeting purified recombinant RBD (0.53 µg rRBD) adjuvanted with poly I:C did not trigger a detectable anti-RBD IgG response, in sharp contrast with Clec9A-RBD immunized mice, which readily produced detectable antibody titers as early as 7 days post immunization **(Figure 1A)**, thereby highlighting the potency of this targeting approach. We next monitored longitudinally the RBD-specific humoral response in Clec9A-RBD immunized mice **(Figure 1B)**. The anti-RBD IgG titers were sustained up to 21 months post-immunization, with a clear T_H_1 isotype profile **(Figure 1C)**. Furthermore, the neutralizing and antibody-dependent cell-mediated cytotoxicity (ADCC) activities **(Figure 1 D&E)**, as well as the binding affinity **(Figure 1F)** of the immune sera against matching SARS-CoV-2 ancestral strain not only persisted throughout the 21 months but progressively increased over time. These observations thus strongly suggested prolonged affinity maturation in Clec9A-RBD immunized mice.

**Figure 1.**
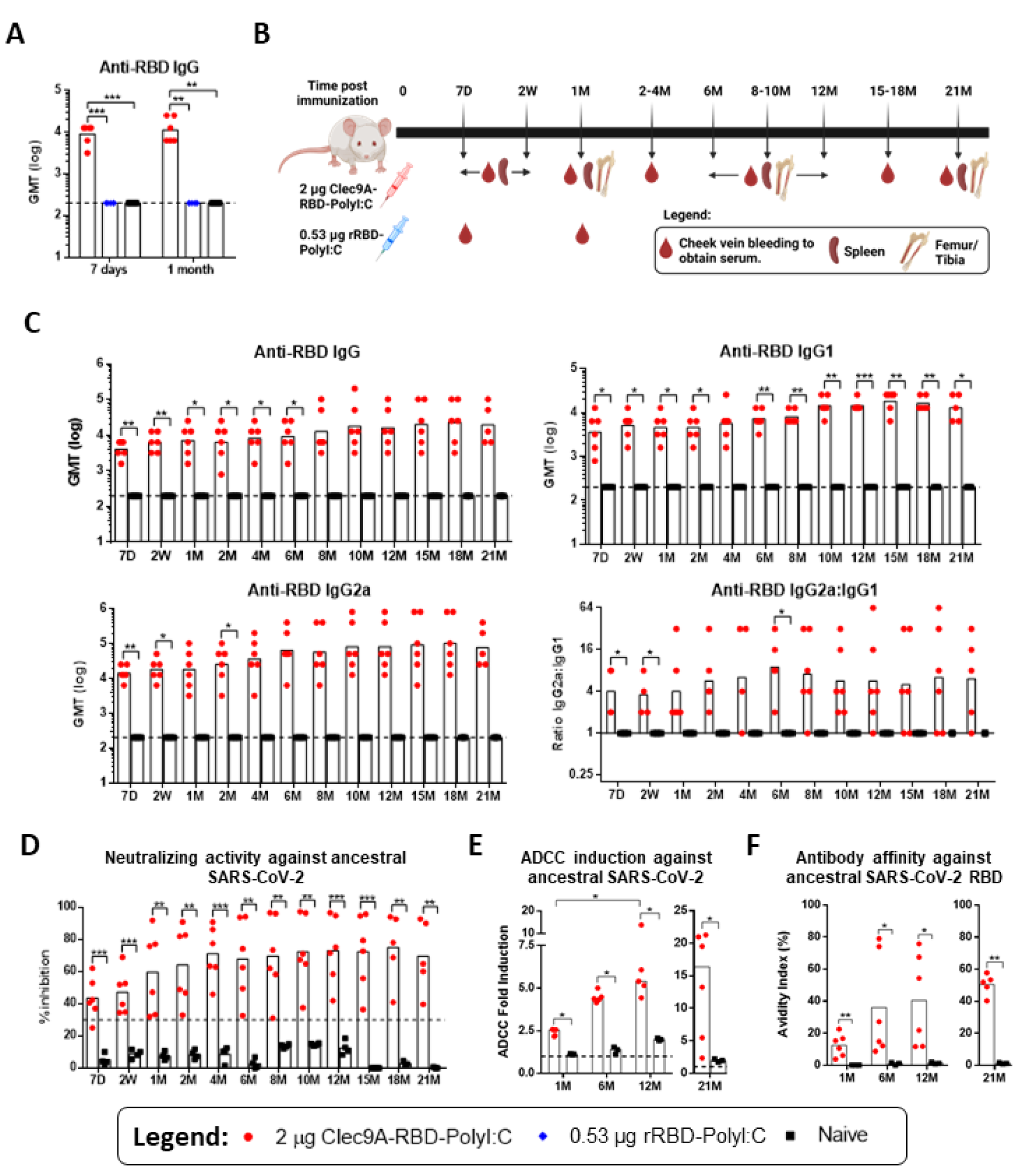
RBD-specific antibody responses induced upon single dose immunization with Clec9A-RBD. **(A)** 5-6 weeks old Balb/c mice (n=5-6 per group) were sc. immunized once with 2 µg of Clec9A-RBD or 0.53 µg rRBD, both adjuvanted with 50 µg Poly I:C. Serum SARS-CoV-2 RBD-specific IgG titers were measured by ELISA at 7 days and 1 month post-immunization. **(B)** Overview of the timeline and immunological assays performed in mice sc. immunized once with 2 µg Clec9A-RBD + 50 µg Poly I:C. **(C)** Longitudinal monitoring of serum SARS-CoV-2 RBD-specific IgG, IgG1, and IgG2a titers by ELISA. The dashed line represents the limit of detection (2.3 log). The ratio of anti-RBD IgG2a: IgG1 was also calculated. **(D)** Longitudinal monitoring of serum neutralizing activity against ancestral SARS-CoV-2 by c-Pass sVNT. The dashed line at 30% represents the cut-off value above which samples are considered positive. **(E)** ADCC activity and **(F)** IgG avidity index of immune sera against ancestral SARS-CoV-2 at 1-, 6-, 12-, and 21 months post-immunization were determined by ADCC Reporter Bioassay, and urea wash ELISA respectively. The dashed line represents the baseline ADCC fold induction (1.0). **(B-F)** Symbols represent individual animals and bars represent **(B, C)** geometric mean, **(D, F)** mean, and **(E)** median. Statistical analysis: **(B)** One-Way ANOVA with Tukey’s correction for multiple comparisons, **(C, D, F)** unpaired two tailed *t* test, and **(E)** Mann-Whitney test (within same timepoint) or two tailed Friedman test with Dunnett’s correction for multiple comparisons (between different timepoints). *p < 0.05, **p < 0.01, or ***p < 0.001.

### Temporal enhancement in the potency and breadth of RBD-specific antibody response against SARS-CoV-2 variants and sarbecoviruses

We next assessed the neutralizing activity of Clec9A-RBD immune sera against a panel of 20 sarbecoviruses, which includes SARS-CoV-2 variants of concern (VOCs), SARS-CoV, and animal CoV from the bat and pangolin lineages [25]. The neutralizing activity against most SARS-CoV-2 VOCs increased over time and persisted throughout the 21-month monitoring period **(Figure 2A and S2A)**, except for Omicron lineages, which was not surprising given the extended repertoire of immune escape mutations and in line with previous reports [26]. Interestingly, neutralizing activity against clade-1B sarbecoviruses from bat and pangolin species (RaTG13, BANAL-52, BANAL-236, GD-1, Gx-P5L) was readily detected at all the time points tested, that increased progressively throughout the 21-month monitoring period **(Figures 2A and S2A)**. In contrast, limited neutralizing activity was seen against clade-1A sarbecoviruses (SARS-CoV, Rs2018B, RsSHC014, LYRa11 and WIV-1) **(Figures 2A and S2A).**

**Figure 2.**
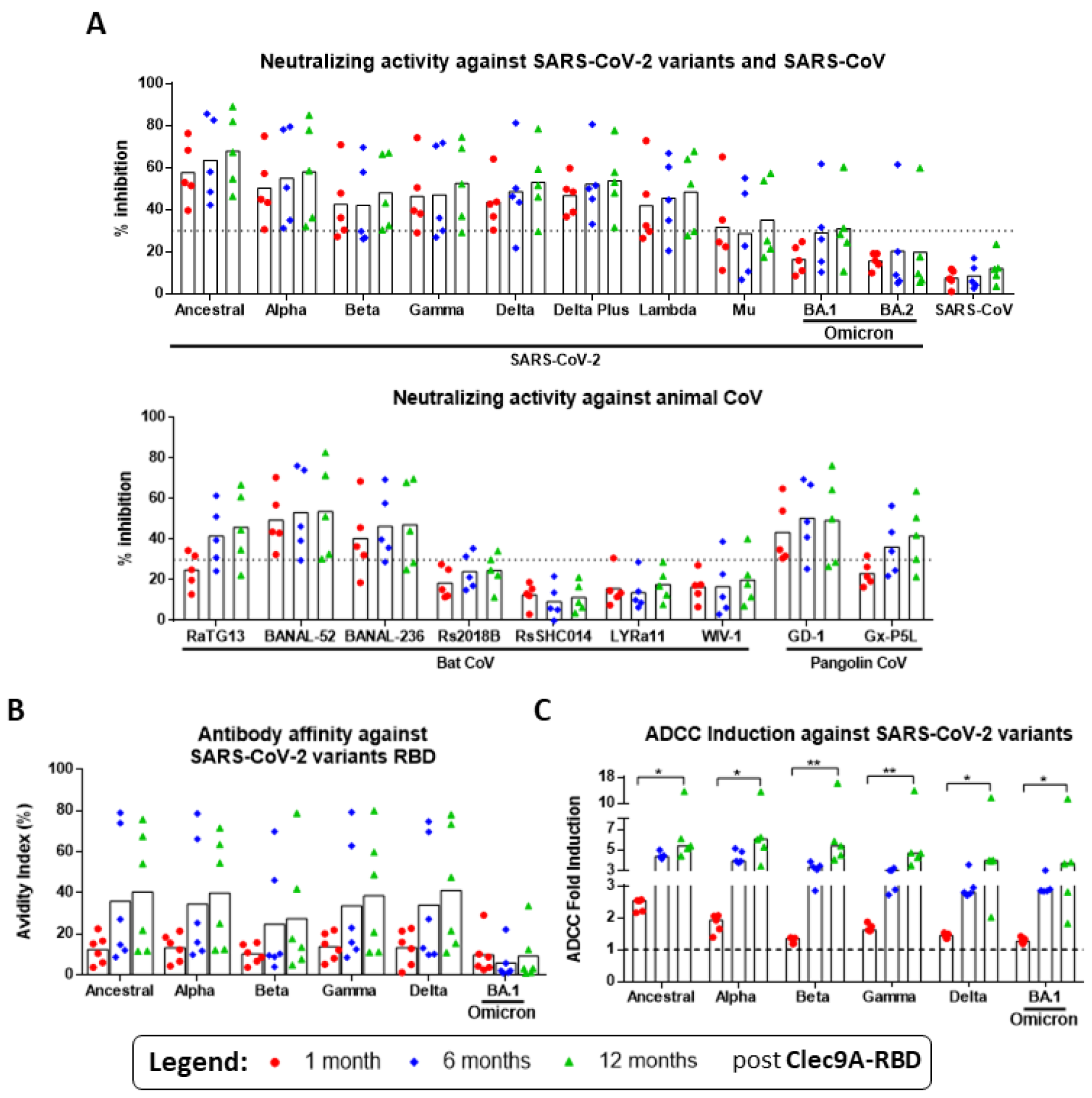
Binding affinity, neutralizing and ADCC activities of immune sera against SARS-CoV-2 variants and other sarbecoviruses. 5-6 weeks old Balb/c mice (n=5 per group) were sc. immunized once with 2 µg of Clec9A-RBD adjuvanted with 50 µg Poly I:C. **(A)** Neutralizing activity of immune sera against a panel of 20 sarbecoviruses at 1-, 6-, and 12 months post-immunization was determined by Multiplex sVNT. The dashed line at 30% represents the cut-off value above which samples are considered positive. **(B)** IgG avidity index and **(C)** ADCC activity of immune sera against ancestral SARS-CoV-2 at 1-, 6-, and 12 months post-immunization were determined by urea wash ELISA, and ADCC Reporter Bioassay, respectively. The dashed line in (C) represents the baseline ADCC fold induction (1.0). **(A-C)** Symbols represent individual animals and bars represent **(B, C)** mean and **(A)** median. Statistical analysis: **(A)** two tailed Friedman test with Dunnett’s correction for multiple comparisons. *p < 0.05 or **p < 0.01.

Binding affinity of the Clec9A-RBD immune sera against SARS-CoV-2 VOCs also increased over time and was maintained even at 21 months post-immunization **(Figures 2B and S2B)**. Similarly, the ADCC activity against SARS-CoV-2 VOCs increased over time and remained high at 21 months post-immunization **(Figure 2C and S2C)**. Excitingly, ADCC activity of the immune sera against Omicron was comparable with that measured against the other VOCs **(Figure 2C and S2C)**, suggesting that non-neutralizing antibodies may contribute to protection against this variant. Together, these results demonstrated that single dose immunization with Clec9A-RBD triggered humoral responses with increasingly broad functional activities over time against a diverse repertoire of sarbecoviruses.

### Sustained RBD-specific T_FH_ cell and B cell subsets after single dose immunization with Clec9A-RBD

The prolonged and increased functional activities of Clec9A-RBD immune sera over time suggested affinity maturation of antibodies, which requires the presence of germinal centres (GC) in immunized mice. We thus quantified RBD-specific T follicular helper (T_FH_) cells and GC B cells. Antigen-specific spleen T_FH_ cell response was generated as early as 7 days post-Clec9A-RBD immunization, which could still be detected 6-12 months later, and was cross-reactive across most SARS-CoV-2 VOCs, including Omicron **(Figure 3A)**. Consistently, GCs were readily detected at 7 days and 2 weeks post-immunization in spleen and brachial lymph nodes respectively, and remarkably, they persisted even after 6-12 months **(Figures 3B and S3)**. RBD-specific GC B cells were also detected in the spleen of immunized mice throughout the monitoring period **(Figure 3C).** Furthermore, RBD-specific memory B cells, plasmablasts and plasma cells were detected in the spleen **(Figure 3D)**; and RBD-reactive antibody secreting cells were found in the bone marrow from Clec9A-RBD immunized mice at 1, 6 and 12 months post-immunization, which were largely cross-reactive against the other VOCs except Omicron **(Figures 3E)**.

**Figure 3.**
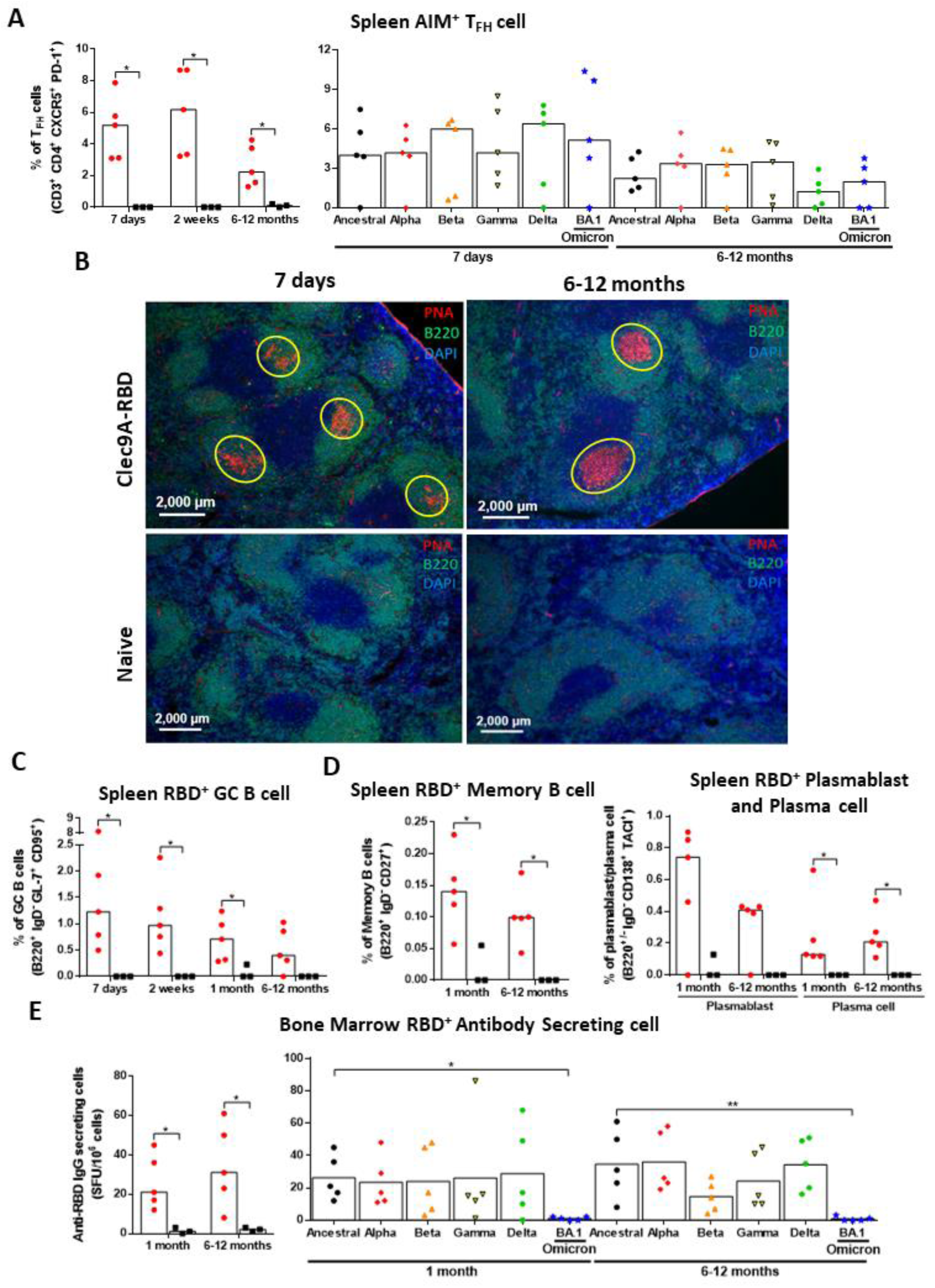
Analysis of RBD-specific T_FH_ and B cell subsets induced upon single dose immunization with Clec9A-RBD. 5-6 weeks old Balb/c mice (n=5 per group) were sc. immunized once with 2 µg of Clec9A-RBD adjuvanted with 50 µg Poly I:C. **(A)** Percentage of AIM^+^ TFH cells (CD3^+^ CD4^+^ CXCR5^+^ PD-1^+^) in response to re-stimulation with ancestral (left panel) and variant (right panel) SARS-CoV-2 RBD peptides in spleens harvested at 7 days, 2 weeks, and 6-12 months (n=2 at 6 and 9 months old each, and n=1 at 12 months old) post-immunization. **(B)** Immune detection of GC B cells in spleen sections from immunized and naive mice harvested at 7 days and 6-12 months post-immunization. GC B cells were defined as B220^+^ PNA^+^ (circled in yellow). DAPI stained cell nuclei. Representative overlay images are shown (scale bar: 2,000 µm). **(C)** Percentage of spleen RBD-specific GC B cells (B220^+^ IgD^-^ GL-7^+^ CD95^+^) measured at 7 days, 2 weeks, 1 month and 6-12 months post-immunization by flow cytometry. **(D)** Percentage of spleen RBD-specific memory B cells (B220^+^ IgD^-^ CD27^+^), plasmablasts (B220^+^ IgD^-^ CD138^+^ TACI^+^) and plasma cells (B220^-^ IgD^-^ CD138^+^ TACI^+^) measured at 1 month and 6-12 months post-immunization by flow cytometry. **(E)** Frequency of bone marrow antibody secreting cells present reactive to ancestral (left panel) and variant (right panel) SARS-CoV-2 RBD determined by B cell ELISPOT. SFU = spot forming unit. **(A, C-E)** Symbols represent individual animals and bars represent the median. Statistical analysis: **(A, C-E)** Mann Whitney test or Kruskal Walis test with Dunnett’s correction for multiple comparisons. *p < 0.05 or **p<0.01.

Together, these data indicated that single dose immunization with Clec9A-RBD induced the formation of GC with antigen-specific T_FH_ cells and GC B cells that could persist up to at least 12 months post-immunization, with concomitant generation of memory B cells, plasmablasts, plasma cells and long-lasting antibody secreting cells. This likely explains the prolonged and sustained affinity maturation observed in the immunized mice.

### Sustained, poly-functional, cross-reactive RBD-specific cellular responses following single dose immunization with Clec9A-RBD

We next investigated the RBD-specific T cell responses upon Clec9A-RBD single dose immunization. Compared to mice immunized with an antigen equivalent amount of purified rRBD, Clec9A-RBD immunized mice produced significantly higher IFN-γ secreting cells upon splenocyte re-stimulation with RBD peptides, highlighting the superior ability of Clec9A-targeting to induce cellular immune responses **(Figure 4A&B)**. In mice immunized with Clec9A-RBD, the relative abundance of T cells producing IL-5 and IFN-γ respectively indicated strong T_H_1-biased cellular responses against matching SARS-CoV-2 strain and VOCs **(Figure 4C)**. It is interesting to note that re-stimulation with Omicron-specific RBD peptides led to responses of a comparable magnitude to the matching strain and the other VOCs, supporting the presence of conserved T cell epitopes in RBD sequence shared amongst the ancestral strain and all the VOCs. Moreover, although most cytokine producing cells were mono-functional (IFN-γ, IL-2, or TNF-α only), Clec9A-RBD single shot immunization elicited a sizeable proportion of poly-functional CD4^+^ and CD8^+^ T cells that secreted either two or three cytokines, and this was observed upon re-stimulation with RBD peptides from the matching strain and all the VOCs, including Omicron **(Figures 4D&E)**. Of note, a greater number of RBD-specific CD8^+^ T cells compared to CD4^+^ T cells was measured in the spleen from immunized mice. Finally and strikingly, IFN-γ, IL-2, and TNF-α secreting cells were detected in spleens harvested at 6-12 months post-immunization and upon re-stimulation with matching or VOCs RBD peptides, demonstrating strong priming of the RBD-specific cellular responses and robust memory T cell generation **(Figure S4)**.

**Figure 4.**
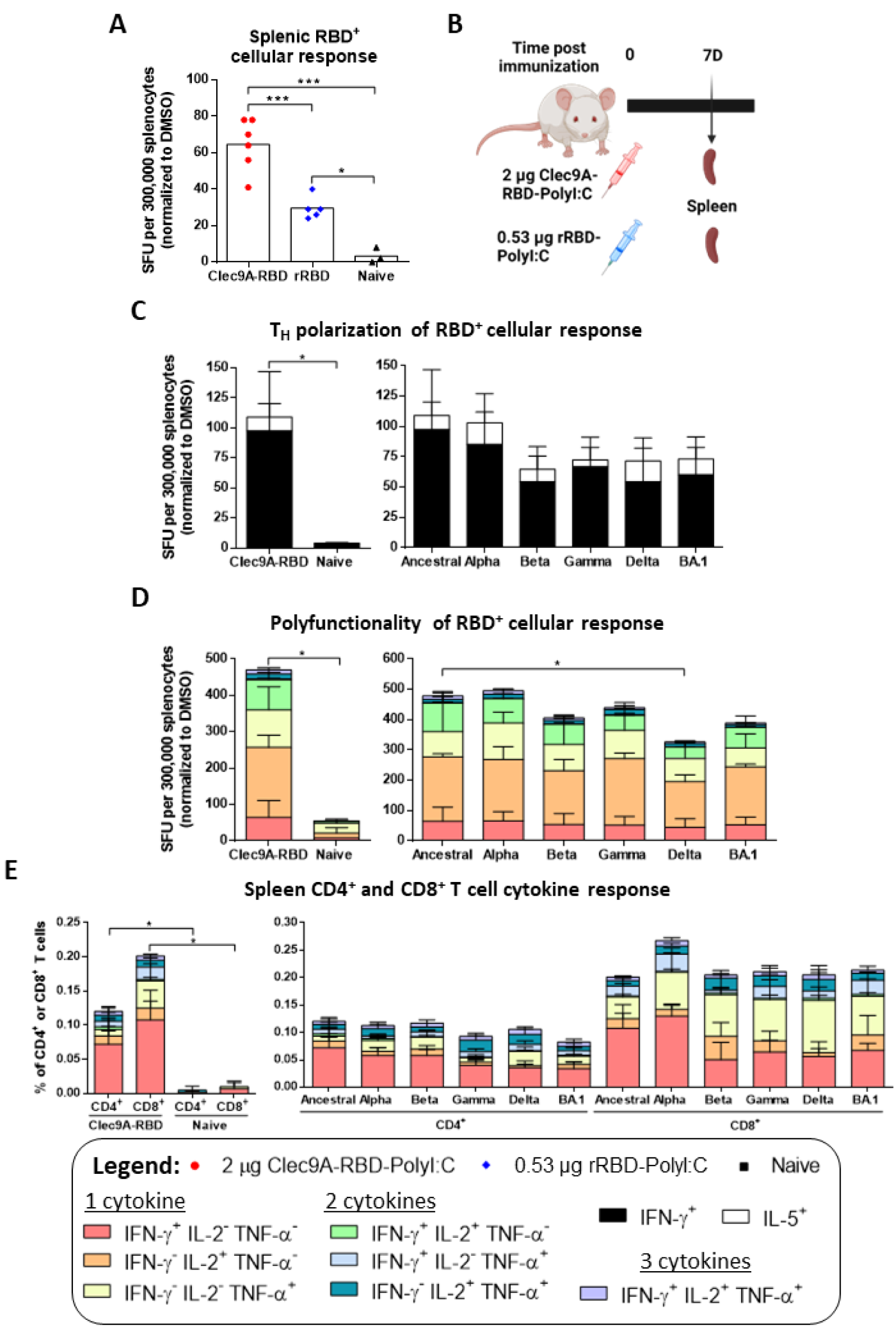
RBD-specific cellular responses induced upon single dose immunization with Clec9A-RBD. **(A)** Frequency of IFN-γ^+^ splenocytes determined by T cell ELISPOT upon re-stimulation with ancestral SARS-CoV-2 RBD peptides 7 days after single dose immunization with 2 µg Clec9A-RBD or 0.53 µg rRBD, both adjuvanted with 50 µg Poly I:C. **(B)** Overview of the immunological assays performed at day 7 post-Clec9A-RBD immunization to analyze the RBD-specific cellular responses. **(C)** Frequency of IFN-γ^+^ and IL-5^+^ splenocytes determined by IFN-γ/IL-5 FluoroSPOT upon re-stimulation with ancestral (left panel) and variant (right panel) SARS-CoV-2 RBD peptides. **(D)** Frequency of splenocytes producing IFN-γ, IL-2, and/or TNF-α determined by IFN-γ/IL-2/TNF-α FluoroSPOT upon re-stimulation with ancestral (left panel) and variant (right) SARS-CoV-2 RBD peptides. **(A, C, D)** SFU = spot forming unit. **(E)** Percentage of spleen CD4^+^ and CD8^+^ T cells expressing IFN-γ, IL-2, and/or TNF-α determined by flow cytometry upon re-stimulation with ancestral (left panel) and variant (right) SARS-CoV-2 RBD peptides. **(A)** Symbols represent individual animals and bars represent the mean **(C-E)** ± standard deviation (SD). Statistical analysis: **(A, C-E)** unpaired two tailed *t* test or One-Way ANOVA with Tukey’s correction for multiple comparisons. *p < 0.05 or ***p < 0.001.

Together, our results indicated that Clec9A-RBD single dose immunization elicited durable poly-functional T_H_1 RBD-specific cellular immune responses that were cross-reactive with all the VOCs tested including Omicron.

### Clec9A-RBD systemic immunization induced RBD-specific humoral and cellular responses in the lungs

Induction of antibody responses and tissue-resident T cells at mucosal sites have been associated with robust protection against SARS-CoV-2 infection in lung tissues [27]. While intranasal vaccines ideally are expected to confer strong mucosal protective immunity, a few systemically administered vaccine candidates have been reported to elicit robust lung mucosal immune responses [28-30]. Hence, we investigated whether systemic single dose immunization with Clec9A-RBD could generate antigen-specific cellular and humoral immunity in the lung tissue **(Figure 5A)**. Impressively, between 2.5 and 4 log_10_ of anti-RBD IgG titers with a strong T_H_1 isotype profile were readily detected in the bronchoalveolar lavage (BAL) fluids harvested two weeks post-Clec9A-RBD immunization **(Figure 5B)**. Moreover, these BAL samples displayed neutralizing and ADCC activity against the matching ancestral SARS-CoV-2 strain **(Figures 5C&D)**. Production of RBD-specific antibodies in the lung was consistent with the presence of RBD-specific GC B cells in this tissue **(Figure 5E)**. Finally, systemic immunization with Clec9A-RBD induced generation of poly-functional RBD-specific CD4^+^ and CD8^+^ T cells in the lung, with again, a greater proportion of CD8^+^ T cells compared to CD4^+^ T cells **(Figure 5F)**.

Collectively, these findings demonstrated that systemic single dose immunization with Clec9A-RBD elicited both humoral and cellular antigen-specific immune responses in the lungs.

**Figure 5.**
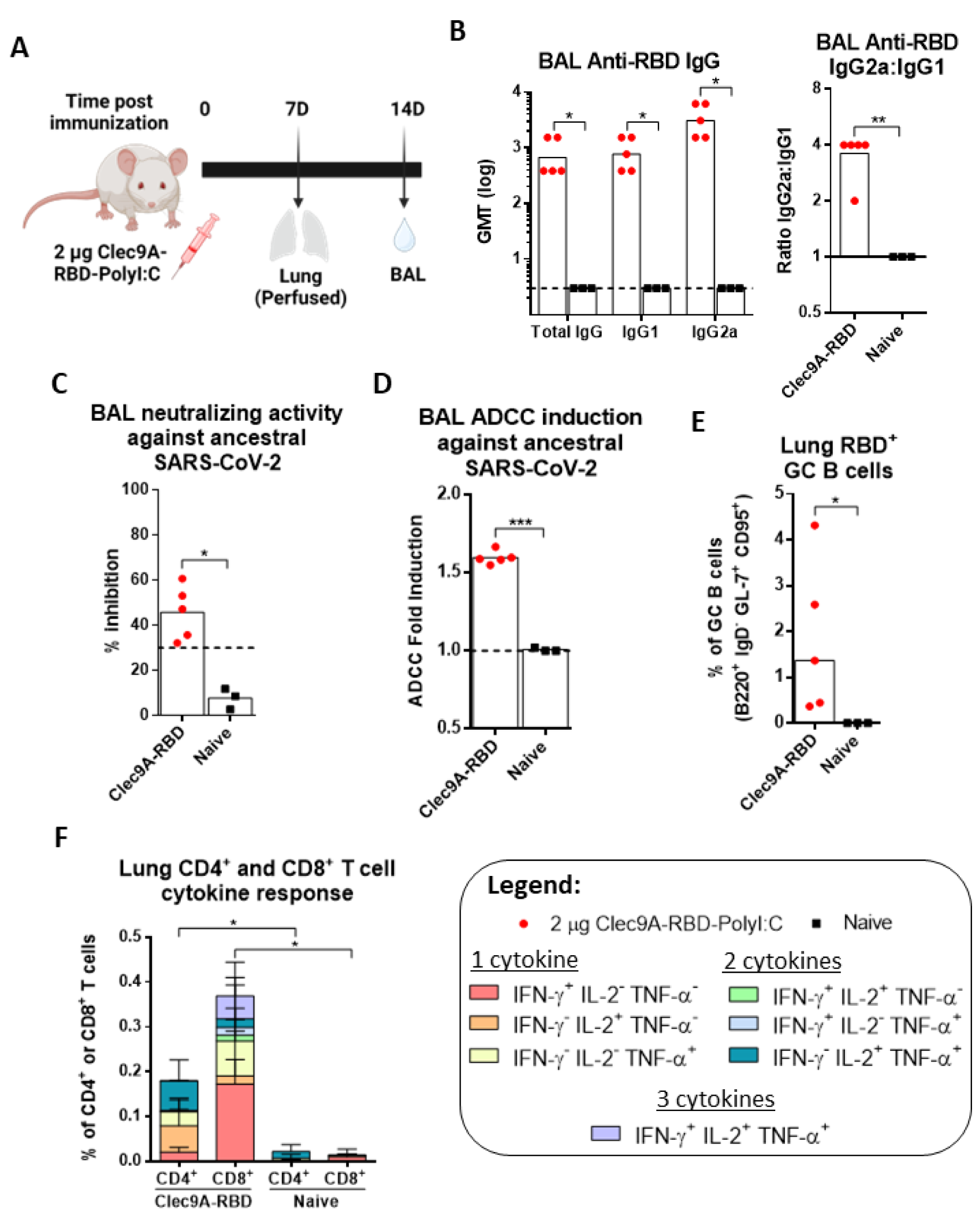
Induction of RBD-specific mucosal immunity upon single dose systemic immunization with Clec9A-RBD. 5-6 weeks old Balb/c mice (n=5 per group) were sc. immunized once with 2 µg Clec9A-RBD adjuvanted with 50 µg Poly I:C. **(A)** Overview of the immunological assays performed to analyze RBD-specific humoral and cellular responses induced in the lungs. **(B)** SARS-CoV-2 RBD-specific IgG, IgG1, and IgG2a titers determined by ELISA in BAL samples harvested at 2 weeks post-immunization. The dashed line represents the limit of detection (0.477 log). The ratio of anti-RBD IgG2a: IgG1 was also calculated (left panel). **(C)** Neutralizing activity against ancestral SARS-CoV-2 determined by c-Pass sVNT in BAL samples harvested at 2 weeks post-immunization. The dashed line at 30% represents the cut-off value above which samples are considered positive. **(D)** ADCC activity against ancestral SARS-CoV-2 determined by ADCC Reporter Bioassay in BAL samples harvested at 2 weeks post-immunization. The dashed line represents the baseline ADCC fold induction (1.0). **(E)** Percentage of RBD-specific GC B cells determined by flow cytometry in perfused lung harvested at 7 days post-immunization and upon re-stimulation with ancestral SARS-CoV-2 RBD peptides. **(F)** Percentage of CD4^+^ and CD8^+^ T cells expressing IFN-γ, IL-2, and/or TNF-α determined by flow cytometry in perfused lung harvested at day 7 post-immunization and upon re-stimulation with ancestral SARS-CoV-2 RBD peptides. **(B-E)** Symbols represent individual animals and bars represent **(B)** geometric mean, **(C-E)** mean and **(F)** mean ± SD. Statistical analysis: **(B-F)** unpaired two tailed *t* test. *p < 0.05 or **p < 0.01.

## DISCUSSION

In this work, we demonstrated that targeting SARS-CoV-2 RBD to Clec9A-expressing DCs (cDC1) resulted in potent and exceptionally sustained immune responses upon single dose immunization. Importantly, we provided evidence of affinity maturation over time, which led to improved potency and breadth of the humoral immune response against a broad representation of the sarbecovirus family. Mechanistically, this can be explained by the strong induction of T_FH_ cells, and persistence of GC reactions which were still detected at 12 months post-immunization. Type-I IFN, IL-6 and IL-12-mediated signaling, coupled with prolonged antigen presentation and CD4^+^ T cell stimulation by DCs and B cells drive the formation and maintenance of T_FH_ cells [23, 31]. As cDC1 are known to secrete or drive the expression of these cytokines, targeting vaccine candidates to this DC subset is hence expected to induce potent T_FH_ responses [14, 20, 32]. Additionally, internalization of Clec9A antibody constructs into cDC1 has been shown to involve the recycling endosomal pathway, which allows the construct to be exposed intact at the cell surface and interact with cognate B cells in close proximity with T_FH_ cells [13, 15, 33, 34], thereby facilitating GC formation and maintenance [31, 35]. As a result, GC reaction and affinity maturation occur, allowing the generation of antigen-specific antibody secreting cells and memory B cells with enhanced functional affinity.

In addition, prolonged antigen availability has also been reported to be essential for GC priming and maintenance [36-38]. Our previous work has shown that Clec9A constructs may persist up to one week in circulation, thereby allowing prolonged antigen presentation [13]. This may therefore further contribute to the persistence of GC and RBD-specific GC B cells up to 12 months post-Clec9A-RBD immunization.

While RBD contains major neutralizing epitopes, it also contains non-neutralizing B cell epitopes that may contribute to protection through non-neutralizing mechanisms such as ADCC, and antibody-dependent cell phagocytosis (ADCP) [39-41]. Consistently, we demonstrated that Clec9A-RBD immune sera displayed significant ADCC activity not only against matching SARS-CoV-2 ancestral strain but also against all the VOCs that were tested including Omicron. The latter observation thus supports the presence of non-neutralizing B cell epitopes in RBD that are more conserved than the neutralizing ones across the VOCs. Therefore, although Omicron successfully escapes neutralizing antibodies raised upon first generation COVID-19 vaccines or infection with an earlier VOC, cross-reactive non-neutralizing antibodies may still confer some level of protection. This is in line with previous studies showing that RBD-specific antibodies were able to protect against lethal SARS-CoV-2 challenge and severe disease through Fc effector mechanisms [39-41].

Our results also showed that Clec9A-RBD single dose immunization induced a potent and sustained antigen-specific poly-functional T_H_1 cellular response, consistent with our previous studies that targeted antigens to Clec9A-expressing cDC1 [16, 42]. Poly-functional T cells have been proposed to represent a correlate of protection in other infectious diseases such as dengue and this feature is applicable towards COVID-19 as well [43-45]. Furthermore, Clec9A-RBD was able to elicit robust RBD-specific T cell memory that could be recalled 6-12 months post-immunization. IL-27 has been found to be critical for the formation and maintenance of memory T cells, whereby cDC1 and B cells were reported to be major producers of this cytokine [46-49]. Since the Clec9A-targeting approach facilitates interaction of T cells with cDC1, coupled with the persistence of RBD-specific B cells, these conditions likely favor IL-27-mediated signaling, thereby promoting induction and maintenance of memory T cells.

Furthermore, we observed that Clec9A-RBD immunization induced prominent CD8^+^ T cell responses despite the fact that SARS-CoV-2 RBD largely comprises of CD4^+^ T cell epitopes, and that RBD-specific cellular responses induced by existing COVID-19 vaccines are CD4^+^ dominated [50-52]. We previously showed that Clec9A targeting resulted in strong CD8^+^ T cell response owing to effective antigen cross-presentation activity in cDC1 [14, 17]. In addition, and importantly, we showed that the T cell responses were cross-reactive against all the VOCs tested, including Omicron. This again is in line with previous work showing that T cell epitopes in spike and RBD are largely conserved and cross-reactive across SARS-CoV-2 variants, and may therefore contribute to protection [12, 52, 53].

Finally, we provided evidence of induction of RBD-specific cellular and humoral responses in the lung tissue following systemic single dose immunization with Clec9A-RBD. Given the prolonged presence of Clec9A-targeting constructs in the circulation, along with Clec9A-expressing CD103^+^ cDC1 residing in the lungs and airway mucosa, it may be possible for the Clec9A-RBD antibody construct to reach the lung tissue for targeted uptake by resident cDC1, allowing effective priming of antigen-specific lung T and B cells [13, 15, 54]. Induction of strong and persistent mucosal immunity represents a key achievement and is expected to confer protection against infection and limit transmission.

In conclusion, we have shown that Clec9A-RBD single dose immunization induces robust and exceptionally sustained and functional humoral and cellular responses, owing to favorable cytokine and cellular microenvironment that promotes and maintains GC reactions and memory T cells. Importantly, the ability to induce both systemic and mucosal (lung) immunity positions Clec9A-RBD as a potential game changer that may limit both infection and transmission. As the world is transitioning towards the endemic phase of COVID-19, the Clec9A targeting technology provides a versatile plug-and-go vaccine platform that can be rapidly updated to address fast evolving variants that circulate in different parts of the world. Importantly, the cDC1 subset equivalent in humans (CD141^+^ DCs) has been found in various tissues and mucosae at constant number throughout the entire human lifespan [55]. Our previous work has shown that Clec9A targeting resulted in potent Ag-specific immune responses in macaques [56] and humanised mice [57]. Hence, translation of this vaccine strategy to human application is a realistic and exciting avenue.

## MATERIALS AND METHODS

### Cell lines

Human embryonic kidney (HEK) epithelial-like cell line HEK293T was maintained in Dulbecco’s Modified Eagle’s Medium (DMEM) (11965-118; GIBCO^™^, Thermo Fisher Scientific, Waltham, Massachusetts, USA) containing 10% Fetal Bovine Serum (FBS) (10270-106; GIBCO^™^, Thermo Fisher Scientific) and 1% Penicillin-Streptomycin (Pen/Strep) (15140122; GIBCO^™^, Thermo Fisher Scientific). Recombinant Jurkat T cells expressing firefly luciferase gene under the control of NFAT response elements with constitutive expression of mouse FcγRIV (mFcγRIV/NFAT-Jurkat) (79733; BPS Bioscience, San Diego, California, USA) were maintained in Roswell Park Memorial Institute (RPMI) 1640 medium (22400-105; GIBCO^™^, Thermo Fisher Scientific) containing 10% FBS, 1% Pen/Strep, 0.5 mg/mL Geneticin^™^ Selective Antibiotic (10131035; GIBCO^™^, Thermo Fisher Scientific), and 2.5 µg/mL Puromycin Dihydrochloride (A1113802; GIBCO^™^, Thermo Fisher Scientific).

### Recombinant proteins, peptides, and plasmids

The Clec9A-RBD construct was generated as described previously [13, 15]. Briefly, rat IgG2a mAb specific against mouse Clec9A (10B4) was genetically fused to one copy of ancestral Wuhan-Hu-1 SARS-CoV-2 RBD (KSFTVEKGIYQTSNFRVQPTESIVRFPNITNLCPFGEVFNATRFASVYAWNRKRISNCVADYSVLYNSASFSTFKCYGVSPTKLNDLCFTNVYADSFVIRGDEVRQIAPGQTGKIADYNYKLPDDFTGCVIAWNSNNLDSKVGGNYNYLYRLFRKSNLKPFERDISTEIYQAGSTPCNGVEGFNCYFPLQSYGFQPTNGVGYQPYRVVVLSFELLHAPATVCGPKKSTNLVKNKCVNFNFNGLTGTG) via an alanine linker at the C-terminal end of each heavy chain. The Clec9A-RBD antibody construct was produced using Freestyle 293F^TM^ cells and 293Fectin^TM^ (ThermoFisher Scientific), purified by affinity chromatography using protein G, and concentration and purity of constructs were validated using absorbance readings at 280nm (Agilent Technologies), Pierce™ BCA Protein Assay Kit (Thermo Fisher Scientific), Bradford protein concentration assay and sodium dodecyl sulfate-polyacrylamide gel electrophoresis (SDS-PAGE) with Coomasie Blue staining respectively.

Purified recombinant SARS-CoV-2 RBD (rRBD) from the ancestral Wuhan-Hu-1 strain and SARS-CoV-2 variants: Alpha (B.1.1.7), Beta (B.1.351), Gamma (P.1), Delta (B.1.617.2), and BA.1 (Omicron), were produced using the Expi293^™^ expression system (A14635; Thermo Fisher Scientific). Ancestral and variant SARS-CoV-2 RBD peptide pools, each comprising of 53 15-mer peptides overlapping by 11 residues, were purchased from Mimotopes Pte Ltd (Clayton, Victoria, Australia) and reconstituted in DMSO (Sigma-Aldrich, St. Louis, Missouri, USA). Plasmids encoding ancestral and variant SARS-CoV-2 spike protein were purchased from Addgene (170442, 170451, 170449, 170450, 172320, 180375; Addgene plasmid, Watertown, Massachusetts, USA).

### Ethics Statement

5-6 weeks old Balb/c mice were purchased from InVivos and housed under specific pathogen-free (SPF) conditions in individual ventilated cages. All described animal experiments were approved by the Institutional Animal Care and Use Committee (IACUC) of NUS under the protocol number R20-0392 and performed in accordance with the guidelines of the National Advisory Committee for Laboratory Animal Research (NACLAR). Animal facilities are AAALAAC-accredited and licensed by the regulatory body Agri-Food and Veterinary Authority of Singapore (AVA). All efforts were made to minimize animal suffering.

### Mice immunization

5-6 weeks old Balb/c mice were subcutaneously (s.c) injected with 2 µg Clec9A-RBD (unless otherwise specified in the figure legend) or 0.53 µg ancestral SARS-CoV-2 rRBD, supplemented with either 50 µg Polyinosinic-polycytidylic acid (Poly(I:C)) (vac-pic; Invivogen, Massachusetts, USA), 50 µg synthetic Class B CpG oligonucleotide (ODN1826) (vac-1826; Invivogen), or unadjuvanted, topped up to a final volume of 150 µL with sterile phosphate buffered saline (PBS).

### Serum and bronchoalveolar lavage (BAL) fluid collection

Mice were sedated with isofluorane, and blood was collected from the cheek vein at indicated timepoints. After clotting for two hours at room temperature (RT), blood was centrifuged at 6,000 rpm for 10 mins at 10°C to harvest serum for heat inactivation at 56°C for 30 mins before storage at −80°C until further processing. BAL was performed by injecting and aspirating 0.5 mL of 1x PBS containing protease inhibitor cocktail (87786; Thermo Fisher Scientific) twice through the mouse trachea. The recovered BAL fluid was centrifuged at 400 g for 5 mins at 4°C to pellet cell debris before storing the supernatant at −80°C until further analysis.

### Preparation of spleen, bone marrow, and lung single cell suspensions

Following euthanasia at indicated timepoints, spleens, femurs, tibias, and lungs (after perfusion with 20 mL 1x PBS injected into the cardiac right ventricle) were harvested from mice. Single cell suspension of spleen was obtained via mechanical disruption in complete RPMI (cRPMI) (RPMI 1640 medium + 10% FBS + 1% Pen/Strep). Femurs and tibias were cut at one end, centrifuged at maximum speed for 1 min to pellet the bone marrow cells, and resuspended in 1 mL cRPMI. Lungs were minced into small pieces and digested with 0.1 mg/mL Liberase (5401020001; Merck Millipore, Burlington, Massachusetts, USA) in RPMI 1640 medium at 37°C and 5% CO_2_ for 45 mins. All cell suspensions were passed through a 70 µm cell strainer and centrifuged at 400 g for 5 mins at 4°C. To remove red blood cells (RBC), 2 mL of in-house 1x RBC lysis buffer (8.26 g NH_4_Cl, 1 g KHCO_3_, 200 µL 0.5 M EDTA, in 1 L dH_2_O) were added to cells for 5 mins at RT. Subsequently, 10 mL of chilled cRPMI was added to quench the reaction, and cells were centrifuged as above. Cells were then resuspended in cRPMI and counted using the Countess 3 FL Automated Cell Counter (Thermo Fisher Scientific).

### Enzyme-Linked Immunosorbent Assay (ELISA)

#### Anti-RBD IgG and IgG isotype titers

96-well ELISA plates (Flat bottom EIA/RIA high binding) (3590; Corning Life Sciences, Corning, New York, USA) were coated with 5 µg/mL (250 ng per 50 µL) ancestral SARS-CoV-2 rRBD per well and incubated overnight at 4°C. Mouse sera were diluted 1:200 in blocking buffer (1x PBS, 5% non-fat dry milk), followed by two-fold serial dilution. ELISA plates were washed thrice with washing buffer (1x PBS, 0.05% Tween-20) before addition of serially diluted mouse sera. Following overnight incubation at 4°C, plates were washed thrice before addition of horseradish peroxidase (HRP)-conjugated anti-mouse IgG (170-6516; Bio-rad, Hercules, California, USA), anti-mouse IgG1 (ab97240; Abcam, Cambridge, UK), and anti-mouse IgG2a (ab97245; Abcam), diluted at 1:10,000 in blocking buffer. After overnight incubation at 4°C, plates were washed thrice and developed by addition of o-phenylenediamine dihydrochloride (OPD) substrate (P9187; Sigma-Aldrich), and incubation at RT for 10 mins in the dark. The reaction was stopped with 1 M concentrated sulfuric acid (H_2_SO_4_), and absorbance was measured at 490 nm using the Sunrise^™^ absorbance microplate reader (Tecan, Mannedorf, Switzerland). RBD-specific antibody titers were determined by non-linear regression as the reciprocal of the highest serum dilution with an absorbance corresponding to three times the absorbance of blank control wells.

#### Anti-RBD IgG Avidity

Two ELISA plates were coated with 2 µg/mL (100 ng per 50 µL) ancestral or variant SARS-CoV-2 rRBD per well and incubated overnight at 4°C. Using a SARS-CoV-2 spike RBD mAb of known concentration (MAB10580; R&D Systems, Minneapolis, Minnesota, USA) as a molecular standard, anti-RBD IgG in mouse sera were quantified and standardized to 1 µg/mL before further dilution in blocking buffer (1:100). Diluted sera were added into the plates after three washes. Following overnight incubation at 4°C, 5.5 M urea (U5128; Sigma-Aldrich) was added to the wells of one plate, while washing buffer was added to the other, and both plates were incubated at RT for 10 mins. The plates were then washed thrice before addition of HRP-conjugated anti-mouse IgG (1:10,000). Following overnight incubation at 4°C, plates were washed thrice and developed by addition of 3,3’,5,5’-Tetramethylbenzidine (TMB) substrate (00-4201-56; Thermo Fisher Scientific), and incubation at RT for 10 mins in the dark. The reaction was stopped with 1 M H_2_SO_4_, and absorbance was measured at 450 nm as described above. IgG avidity index was calculated as a measurement of antibody functional affinity using the following formula: A_450 nm_ (urea) / A4_50 nm_ (no urea) x 100%, normalized to their respective blanks.

### Surrogate Virus Neutralization Test (sVNT)

#### cPass^™^ sVNT

The cPass^™^ sVNT (L00847-A; GenScript, Piscataway, New Jersey, USA) was used to measure serum and BAL fluid neutralizing antibodies against ancestral SARS-CoV-2, whereby procedure and analysis of neutralizing activity were performed as per manufacturer’s protocol. Serum and BAL fluid samples were diluted 1:10 or used neat respectively.

#### Multiplex sVNT

Multiplex sVNT was established using the Luminex platform as described previously [25]. Briefly, RBDs from 20 sarbecoviruses (**Table S1**) were enzymatically biotinylated, and coated on MagPlex-Avidin microspheres (Luminex, Austin, Texas, USA) at 5 µg per 10^6^ beads. 25 µL of RBD-coated beads (600 beads per antigen) were preincubated with equal volume of mouse serum at a final dilution of 1:20 for 15 mins at 37°C with 200 rpm agitation. Subsequently, 50 µL of phycoerythrin (PE)-conjugated human ACE2 (Genscript) was added to each well and incubated for an additional 15 mins at 37°C with agitation. After two washes with 1% BSA in PBS, the final readings were acquired using the MAGPIX system (Luminex, xPONENT 4.2) following the manufacturer’s instructions. Neutralizing activity was assessed and reported as percentage inhibition using the following formula: [(mean fluorescence intensity (MFI) of 30 negative pre-pandemic samples – individual sample FI) / MFI of 30 negative pre-pandemic samples] x 100%, and a cut-off of ≥ 30% inhibition was defined as positive for neutralizing antibodies against the sarbecovirus.

### Antibody-dependent cell-mediated cytotoxicity (ADCC) Reporter Bioassay

The mouse FcγRIV ADCC Reporter Bioassay was used to evaluate mouse sera and BAL fluids ADCC activity. HEK293T cells were seeded into a 6-well plate (140685; Thermo Fisher Scientific) at 10^6^ cells per well and incubated overnight at 37°C and 5% CO_2_. The cells were then transfected with ancestral or variant SARS-CoV-2 spike protein encoding plasmid using Lipofectamine 3000 Transfection Reagent (L3000001; Thermo Fisher Scientific). After 24 hours, the transfected cells were harvested and stained with Alexa Fluor 647-anti-SARS-CoV-2 RBD antibody (51-6490-82; Thermo Fisher Scientific). Ancestral and variant spike expressing HEK293T cells were sorted for as target cells using the FACSAria^™^ Fusion Cell Sorter (BD Biosciences, Franklin Lakes, New Jersey, USA). Target cells (1.5×10^4^ in 25 µL per well) of each variant were seeded in serum free RPMI into a 96-well V-bottom plate (249952; Thermo Fisher Scientific). Heat inactivated mouse sera and BAL fluid samples were diluted 1:2 in serum free RPMI or used neat respectively, where 25 µL was added to each well and incubated for 1 hour at 37°C. Incubated target cells were washed with 1x PBS once, and 1.2×10^5^ mFcγRIV/NFAT-Jurkat effector cells (in 75 µL serum free RPMI) were then added to each well. The mixture was incubated at 37°C and 5% CO_2_ for 6 hours. Subsequently, 60 µL of cell mixture was transferred into a 96-well white flat-bottom plate (3917; Corning Life Sciences), and an equal volume of Bio-Glo^™^ Luciferase reagent (G7940; Promega, Madison, Wisconsin, USA) was added to each well and rocked gently at RT for 15 mins. Relative luminescence unit (RLU) was measured using the SpectraMax iD3 microplate reader (Molecular Devices, San Jose, California, USA), and ADCC fold induction was calculated as: (RLU sample – RLU blank) / (RLU no serum – RLU blank).

### Enzyme-Linked Immunosorbent Spot (ELISPOT)

#### T cell ELISPOT

RBD-specific T cell responses were investigated using the mouse IFN-γ ELISPOT Set (551083; BD Biosciences), as per manufacturer’s protocol. Splenocytes (3×10^5^ per well) were stimulated overnight with ancestral SARS-CoV-2 RBD peptides at 2 µg/mL diluted in cRPMI. DMSO (34869; Sigma-Aldrich) and 50 ng/mL PMA (tlrl-pma; Invivogen) with 1 µg/mL Ionomycin (inh-ion; Invivogen) were also included as negative and positive controls respectively. Spots were enumerated using the Mabtech IRIS^™^ ELISPOT and FluoroSPOT reader (Mabtech, Nacka Strand, Sweden). Analysis was performed using Mabtech Apex^™^ software version 1.1.5.74, and data was expressed as spot forming units (SFU) per 3×10^5^ splenocytes.

#### B cell ELISPOT

RBD-specific bone marrow antibody secreting cells were detected via B cell ELISPOT. Here, MultiScreen-IP Filter plates (MAIPSWU10; Merck Millipore) were pre-wetted with 50 µL of 70% ethanol for ≤ 2 mins, washed thrice with sterile 1x PBS, and coated with 100 µL per well of 10µg/mL ancestral or variant rRBD, diluted in 1x PBS. Following overnight incubation at 4°C, the plates were washed thrice with sterile 1x PBS and blocked with 200 µL per well of cRPMI for two hours at RT. The blocking solution was discarded and 200 µL of 10^6^ bone marrow cells were added to each well. After incubation at 37°C and 5% CO_2_ for 6 hours, the plates were washed thrice with washing buffer, and 1.5 µg/mL of HRP-conjugated anti-mouse IgG (62-6520; Thermo Fisher Scientific) diluted in 1x PBS was added at 100 µL per well and incubated overnight at 4°C. The plates were then washed thrice with washing buffer, and twice with 1x PBS. 100 µL of AEC substrate (551951; BD Biosciences) was added per well and incubated for 30 mins before rinsing ten times with dH_2_O. Spots were enumerated and analyzed as above, and data was expressed as SFU per 10^6^ bone marrow cells.

### FluoroSPOT

To assess T-helper phenotype and poly-functionality of RBD-specific cellular response, mouse IFN-γ/IL-5 (X-41A43B; Mabtech) and IFN-γ/IL-2/TNF-α (X-41A42B45W; Mabtech) FluoroSPOT were respectively performed, as per manufacturer’s protocol. Splenocytes (3×10^5^ per well) were stimulated for 36 hours with 2 µg/mL ancestral and variant SARS-CoV-2 RBD peptides with 0.2 µg/mL kit-provided anti-mouse CD28 at 37°C and 5% CO_2_. DMSO and 50 ng/mL PMA with 1 µg/mL Ionomycin were also included as negative and positive controls respectively. Enumeration and analysis of spots were performed similarly as ELISPOT, and data was expressed as SFU per 3×10^5^ splenocytes.

### Flow Cytometry

#### Intracellular cytokine staining

Splenocytes or lung cells (10^6^) were stimulated in a 96-well round-bottom plate with 5 µg/mL ancestral and variant SARS-CoV-2 RBD peptides and 1 µg/mL anti-CD28 (40-0281-M001; Tonbo Biosciences, San Diego, California, USA) at 37°C and 5% CO_2_. DMSO and 50 ng/mL PMA with 1 µg/mL Ionomycin were included as negative and positive controls respectively. After 1 hour, 5 µg/mL brefeldin A (420601; Biolegend, San Diego, California, USA) and 2 µM monensin (420701; Biolegend) were added to each well, and stimulation was allowed to proceed for 5 more hours. Thereafter, cells were washed with fluorescence-activated cell sorting (FACS) staining buffer (1x PBS, 2% FBS, 1 mM EDTA) and blocked with anti-CD16/32 antibody (1:200 in FACS buffer, 553142; BD Biosciences) for 10 mins at RT. Cells were then stained with eFluor^™^ 780 Fixable Viability Dye (1:1,000 in 1x PBS, 65-0865-14; Thermo Fisher Scientific) for 10 mins at 4°C in the dark. Subsequently, cells were washed and stained with surface marker antibodies for 30 mins at 4°C in the dark **(Table S2)**. Cells were then washed and resuspended in Transcription Factor Fix/Perm solution prepared as per manufacturer’s protocol (TNB-0607-KIT; Tonbo Biosciences). Following 30 mins incubation at RT, cells were washed twice with 1x Flow Cytometry Perm Buffer prepared as per manufacturer’s instructions (TNB-0607-KIT; Tonbo Biosciences) before staining with intracellular marker antibodies for 30 mins at 4°C in the dark **(Table S2)**.

#### Activation-induced markers positive (AIM^+^) T_FH_ cells

Splenocytes (10^6^) were stimulated overnight in a 96-well round-bottom plate with 5 µg/mL ancestral and variant SARS-CoV-2 RBD peptides with 1 µg/mL anti-CD28 at 37°C and 5% CO_2_. DMSO and 50 ng/mL PMA with 1 µg/mL Ionomycin were included as negative and positive controls respectively. Subsequently, splenocytes were washed, blocked with anti-CD16/32 antibody, and stained with eFluor^™^ 780 Fixable Viability Dye as above. Thereafter, cells were washed and stained as above with surface marker antibodies **(Table S2)**.

#### RBD-specific B cell subsets

Antigen-specific B cell subsets were detected using biotinylated SARS-CoV-2 RBD protein (793906; Biolegend) in combination with streptavidin (SAv)-fluorophore conjugates as described [7]. Briefly, biotinylated SARS-CoV-2 RBD was multimerised with SAv-BV421 and SAv-PE (563259, 554061; BD Biosciences) individually, at a mass ratio of ∼2:1, in increments of five .volumes per 30 mins. SAv-Alexa Fluor 488 (S11223; Thermo Fisher Scientific) was used as a decoy probe without biotinylated RBD to gate out cells that bind to non-specific domains of the antigen probe apart from RBD. Following the multimerization, RBD-BV421, RBD-PE, and SAv-Alexa Fluor 488 were mixed together with 5 µM free D-biotin (B20656; Thermo Fisher Scientific) to minimize cross-reactivity between probes. Freshly isolated splenocytes (5×10^6^) were blocked with anti-CD16/32 antibody and stained with eFluor^™^ 780 Fixable Viability Dye similarly as above. Cells were then washed and stained with the RBD probe master mix for 1 hour at 4°C in the dark. Subsequently, splenocytes were washed and stained as above with surface marker antibodies corresponding to each B cell subset **(Table S2)**.

#### Data acquisition and analysis

After the final staining step, cells were washed and resuspended in FACS buffer prior to data acquisition on LSRFortessa X-20 analyzer (BD Biosciences). UltraComp eBeads (01-2222-42; Thermo Fisher Scientific) were used for single colour compensation controls. Flow cytometry analysis was conducted using FlowJo (v.10.8.1). Intracellular cytokine staining, AIM^+^ T_FH_ cells, and RBD-specific B cell subsets were gated as indicated (**Figures S5 and S6**).

### Immunofluorescence microscopy

Fresh cryo-sections from mouse spleens and brachial lymph nodes (10 µm) were fixed in acetone, blocked with 0.2% BSA, and stained with rat anti-mouse B220/CD45R (14-0452-82; Thermo Fisher Scientific) (1:200) and biotinylated Peanut Agglutinin (PNA) (B-1075; Vector Labs, Newark, California, USA) (1:800) for 3 hours at RT. Cy2-conjugated donkey anti-rat IgG (712-225-153; Jackson ImmunoResearch, West Grove, Pennsylvania, USA) (1:300) and Streptavidin-Cy3 (016-600-084; Jackson ImmunoResearch) (1:500) were used for detection of B220/CD45R and PNA respectively for 2 hours at RT. Sections were counterstained with 4,6-diamidino-2-phenylindole (DAPI) (D1306; Thermo Fisher Scientific) for cell nuclei visualization and mounted for analysis. A fluorescence widefield microscope (Axio Imager.Z1, Axioxam HRM camera; Carl Zeiss Micro Imaging, Inc., Jena, Germany) was used to visualize and image the specimens.

### Statistical analysis

All experiments were conducted independently once or twice, and all *in vitro* and *ex vivo* assays were performed in technical duplicates. Details about the group size, descriptive statistics reported, and statistical tests used, are provided in each corresponding figure legend. All results and data were analyzed using PRISM GraphPad Version 7, and statistical significance is indicated by *p<0.05, **p<0.01, and ***p<0.001.

## List of Supplementary Materials

-Fig S1 to S6

-Tables S1 and S2

## ACKNOWLEDGEMENTS

We would like to thank the FACS and histology core facilities at Life Sciences Institute, National University of Singapore to facilitate sample processing and analyses. We would also like to thank Prof Kim Good-Jacobson from Monash University for insightful discussions.

## Author Contributions

Conceptualization: NYZC, MHL, SA

Performing experiments: NYZC, PST, KP, WCY, BYLC, KST, KMT, AYYY, CHQT, XQ, DJWT

Reagents, inputs: IC, YJT, PAM, CWT, MHL Data analysis: NYZC, SA

Funding acquisition: SA, MHL Supervision: SA

Writing – original draft: NYZC Writing – review & editing: SA, MHL

## Competing interests

MHL, IC and KMT are listed as inventors on patents relating to Clec9A antibodies.

All the authors declare that they have no conflict of interest.

**Data and materials availability:** Anti-Clec9A antibodies are available under material transfer agreement or research collaborative agreement.

## REFERENCES

1. World Health Organization (WHO). (2022). WHO Coronavirus Disease (COVID-19) Dashboard.

2. Neerukonda, S. N., & Katneni, U. (2020). A Review on SARS-CoV-2 Virology, Pathophysiology, Animal Models, and Anti-Viral Interventions. Pathogens (Basel, Switzerland), 9(6), 426.

3. Yue, H., Bai, X., Wang, J., Yu, Q., Liu, W., Pu, J., Wang, X., Hu, J., Xu, D., Li, X., Kang, N., Li, L., Lu, W., Feng, T., Ding, L., Li, X., Qi, X., & Gansu Provincial Medical Treatment Expert Group of COVID-19 (2020). Clinical characteristics of coronavirus disease 2019 in Gansu province, China. Annals of palliative medicine, 9(4), 1404–1412.

4. Zhou, F., Yu, T., Du, R., Fan, G., Liu, Y., Liu, Z., Xiang, J., Wang, Y., Song, B., Gu, X., Guan, L., Wei, Y., Li, H., Wu, X., Xu, J., Tu, S., Zhang, Y., Chen, H., & Cao, B. (2020). Clinical course and risk factors for mortality of adult inpatients with COVID-19 in Wuhan, China: a retrospective cohort study. Lancet (London, England), 395(10229), 1054–1062.

5. Mehandru, S., & Merad, M. (2022). Pathological sequelae of long-haul COVID. Nature immunology, 23(2), 194–202.

6. Hwang, J. K., Zhang, T., Wang, A. Z., & Li, Z. (2021). COVID-19 vaccines for patients with cancer: benefits likely outweigh risks. Journal of hematology & oncology, 14(1), 38.

7. Goel, R. R., Painter, M. M., Apostolidis, S. A., Mathew, D., Meng, W., Rosenfeld, A. M., Lundgreen, K. A., Reynaldi, A., Khoury, D. S., Pattekar, A., Gouma, S., Kuri-Cervantes, L., Hicks, P., Dysinger, S., Hicks, A., Sharma, H., Herring, S., Korte, S., Baxter, A. E., Oldridge, D. A., … Wherry, E. J. (2021). mRNA vaccines induce durable immune memory to SARS-CoV-2 and variants of concern. Science (New York, N.Y.), 374(6572), abm0829.

8. Vardhana, S., Baldo, L., Morice, W. G., 2nd, & Wherry, E. J. (2022). Understanding T cell responses to COVID-19 is essential for informing public health strategies. Science immunology, 7(71), eabo1303.

9. Chemaitelly, H., Ayoub, H. H., Tang, P., Hasan, M. R., Coyle, P., Yassine, H. M., Al-Khatib, H. A., Smatti, M. K., Al-Kanaani, Z., Al-Kuwari, E., Jeremijenko, A., Kaleeckal, A. H., Latif, A. N., Shaik, R. M., Abdul-Rahim, H. F., Nasrallah, G. K., Al-Kuwari, M. G., Butt, A. A., Al-Romaihi, H. E., Al-Thani, M. H., … Abu-Raddad, L. J. (2022). Immune Imprinting and Protection against Repeat Reinfection with SARS-CoV-2. The New England journal of medicine, 10.1056/NEJMc2211055. Advance online publication.

10. Park, Y.-J., Pinto, D., Walls, A. C., Liu, Z., De Marco, A., Benigni, F., Zatta, F., Silacci-Fregni, C., Bassi, J., Sprouse, K. R., Addetia, A., Bowen, J. E., Stewart, C., Giurdanella, M., Saliba, C., Guarino, B., Schmid, M. A., Franko, N. M., Logue, J. K., … Veesler, D. (2022). Imprinted antibody responses against SARS-COV-2 omicron sublineages. Science, 378(6618).

11. Mouro, V., & Fischer, A. (2022). Dealing with a mucosal viral pandemic: lessons from COVID-19 vaccines. Mucosal immunology, 15(4), 584–594.

12. Macri, C., Dumont, C., Johnston, A. P., & Mintern, J. D. (2016). Targeting dendritic cells: a promising strategy to improve vaccine effectiveness. Clinical & translational immunology, 5(3), e66.

13. Lahoud, M. H., Ahmet, F., Kitsoulis, S., Wan, S. S., Vremec, D., Lee, C. N., Phipson, B., Shi, W., Smyth, G. K., Lew, A. M., Kato, Y., Mueller, S. N., Davey, G. M., Heath, W. R., Shortman, K., & Caminschi, I. (2011). Targeting antigen to mouse dendritic cells via Clec9A induces potent CD4 T cell responses biased toward a follicular helper phenotype. Journal of immunology (Baltimore, Md.: 1950), 187(2), 842–850.

14. Tullett, K. M., Lahoud, M. H., & Radford, K. J. (2014). Harnessing Human Cross-Presenting CLEC9A (+) XCR1(+) Dendritic Cells for Immunotherapy. Frontiers in immunology, 5, 239.

15. Park, H. Y., Tan, P. S., Kavishna, R., Ker, A., Lu, J., Chan, C., Hanson, B. J., MacAry, P. A., Caminschi, I., Shortman, K., Alonso, S., & Lahoud, M. H. (2017). Enhancing vaccine antibody responses by targeting Clec9A on dendritic cells. NPJ vaccines, 2, 31.

16. Kavishna, R., Kang, T. Y., Vacca, M., Chua, B., Park, H. Y., Tan, P. S., Chow, V. T., Lahoud, M. H., & Alonso, S. (2022). A single-shot vaccine approach for the universal influenza A vaccine candidate M2e. Proceedings of the National Academy of Sciences of the United States of America, 119(13), e2025607119.

17. Lahoud, M. H., & Radford, K. J. (2022). Enhancing the immunogenicity of cancer vaccines by harnessing CLEC9A. Human vaccines & immunotherapeutics, 18(1), 1873056.

18. Zhang, J. G., Czabotar, P. E., Policheni, A. N., Caminschi, I., Wan, S. S., Kitsoulis, S., Tullett, K. M., Robin, A. Y., Brammananth, R., van Delft, M. F., Lu, J., O’Reilly, L. A., Josefsson, E. C., Kile, B. T., Chin, W. J., Mintern, J. D., Olshina, M. A., Wong, W., Baum, J., Wright, M. D., … Lahoud, M. H. (2012). The dendritic cell receptor Clec9A binds damaged cells via exposed actin filaments. Immunity, 36(4), 646–657.

19. Ahrens, S., Zelenay, S., Sancho, D., Hanč, P., Kjær, S., Feest, C., Fletcher, G., Durkin, C., Postigo, A., Skehel, M., Batista, F., Thompson, B., Way, M., Reis e Sousa, C., & Schulz, O. (2012). F-actin is an evolutionarily conserved damage-associated molecular pattern recognized by DNGR-1, a receptor for dead cells. Immunity, 36(4), 635–645.

20. Ashour, D., Arampatzi, P., Pavlovic, V., Förstner, K. U., Kaisho, T., Beilhack, A., Erhard, F., & Lutz, M. B. (2020). IL-12 from endogenous cDC1, and not vaccine DC, is required for Th1 induction. JCI insight, 5(10), e135143.

21. Ebenig, A., Muraleedharan, S., Kazmierski, J., Todt, D., Auste, A., Anzaghe, M., Gömer, A., Postmus, D., Gogesch, P., Niles, M., Plesker, R., Miskey, C., Gellhorn Serra, M., Breithaupt, A., Hörner, C., Kruip, C., Ehmann, R., Ivics, Z., Waibler, Z., Pfaender, S., … Mühlebach, M. D. (2022). Vaccine-associated enhanced respiratory pathology in COVID-19 hamsters after TH2-biased immunization. Cell reports, 40(7), 111214.

22. Kato, Y., Zaid, A., Davey, G. M., Mueller, S. N., Nutt, S. L., Zotos, D., Tarlinton, D. M., Shortman, K., Lahoud, M. H., Heath, W. R., & Caminschi, I. (2015). Targeting Antigen to Clec9A Primes Follicular Th Cell Memory Responses Capable of Robust Recall. Journal of immunology (Baltimore, Md.: 1950), 195(3), 1006–1014.

23. Krishnaswamy, J. K., Alsén, S., Yrlid, U., Eisenbarth, S. C., & Williams, A. (2018). Determination of T Follicular Helper Cell Fate by Dendritic Cells. Frontiers in immunology, 9, 2169.

24. Wheatley, A. K., Fox, A., Tan, H. X., Juno, J. A., Davenport, M. P., Subbarao, K., & Kent, S. J. (2021). Immune imprinting and SARS-CoV-2 vaccine design. Trends in immunology, 42(11), 956–959.

25. Tan, C. W., Chia, W. N., Zhu, F., Young, B. E., Chantasrisawad, N., Hwa, S. H., Yeoh, A. Y., Lim, B. L., Yap, W. C., Pada, S. K. M. S., Tan, S. Y., Jantarabenjakul, W., Toh, L. K., Chen, S., Zhang, J., Mah, Y. Y., Chen, V. C., Chen, M. I., Wacharapluesadee, S., Sigal, A., … Wang, L. F. (2022). SARS-CoV-2 Omicron variant emerged under immune selection. Nature microbiology, 7(11), 1756–1761.

26. Jacobsen, H., Strengert, M., Maaß, H., Ynga Durand, M. A., Katzmarzyk, M., Kessel, B., Harries, M., Rand, U., Abassi, L., Kim, Y., Lüddecke, T., Metzdorf, K., Hernandez, P., Ortmann, J., Heise, J. K., Castell, S., Gornyk, D., Glöckner, S., Melhorn, V., Kemmling, Y., … Cicin-Sain, L. (2022). Diminished neutralization responses towards SARS-CoV-2 Omicron VoC after mRNA or vector-based COVID-19 vaccinations. Scientific reports, 12(1), 19858.

27. Lavelle, E. C., & Ward, R. W. (2022). Mucosal vaccines - fortifying the frontiers. Nature reviews. Immunology, 22(4), 236–250.

28. Del Campo, J., Pizzorno, A., Djebali, S., Bouley, J., Haller, M., Pérez-Vargas, J., Lina, B., Boivin, G., Hamelin, M. E., Nicolas, F., Le Vert, A., Leverrier, Y., Rosa-Calatrava, M., Marvel, J., & Hill, F. (2019). OVX836 a recombinant nucleoprotein vaccine inducing cellular responses and protective efficacy against multiple influenza A subtypes. NPJ vaccines, 4, 4.

29. Laczkó, D., Hogan, M. J., Toulmin, S. A., Hicks, P., Lederer, K., Gaudette, B. T., Castaño, D., Amanat, F., Muramatsu, H., Oguin, T. H., 3rd, Ojha, A., Zhang, L., Mu, Z., Parks, R., Manzoni, T. B., Roper, B., Strohmeier, S., Tombácz, I., Arwood, L., Nachbagauer, R., … Pardi, N. (2020). A Single Immunization with Nucleoside-Modified mRNA Vaccines Elicits Strong Cellular and Humoral Immune Responses against SARS-CoV-2 in Mice. Immunity, 53(4), 724–732.e7.

31. Steinbuck, M. P., Seenappa, L. M., Jakubowski, A., McNeil, L. K., Haqq, C. M., & DeMuth, P. C. (2021). A lymph node-targeted Amphiphile vaccine induces potent cellular and humoral immunity to SARS-CoV-2. Science advances, 7(6), eabe5819.

32. Webb, L. M. C., & Linterman, M. A. (2017). Signals that drive T follicular helper cell formation. Immunology, 152(2), 185–194.

33. Arabpour, M., Lebrero-Fernandez, C., Schön, K., Strömberg, A., Börjesson, V., Lahl, K., Ballegeer, M., Saelens, X., Angeletti, D., Agace, W., & Lycke, N. (2022). ADP-ribosylating adjuvant reveals plasticity in cDC1 cells that drive mucosal Th17 cell development and protection against influenza virus infection. Mucosal immunology, 15(4), 745–761.

34. Tullett, K. M., Tan, P. S., Park, H. Y., Schittenhelm, R. B., Michael, N., Li, R., Policheni, A. N., Gruber, E., Huang, C., Fulcher, A. J., Danne, J. C., Czabotar, P. E., Wakim, L. M., Mintern, J. D., Ramm, G., Radford, K. J., Caminschi, I., O’Keeffe, M., Villadangos, J. A., Wright, M. D., … Lahoud, M. H. (2020). RNF41 regulates the damage recognition receptor Clec9A and antigen cross-presentation in mouse dendritic cells. eLife, 9, e63452.

35. Kato, Y., Steiner, T. M., Park, H. Y., Hitchcock, R. O., Zaid, A., Hor, J. L., Devi, S., Davey, G. M., Vremec, D., Tullett, K. M., Tan, P. S., Ahmet, F., Mueller, S. N., Alonso, S., Tarlinton, D. M., Ploegh, H. L., Kaisho, T., Beattie, L., Manton, J. H., Fernandez-Ruiz, D., … Caminschi, I. (2020). Display of Native Antigen on cDC1 That Have Spatial Access to Both T and B Cells Underlies Efficient Humoral Vaccination. Journal of immunology (Baltimore, Md.: 1950), 205(7), 1842–1856.

36. Kwun, J., Manook, M., Page, E., Burghuber, C., Hong, J., & Knechtle, S. J. (2017). Crosstalk Between T and B Cells in the Germinal Center After Transplantation. Transplantation, 101(4), 704–712.

37. Lee, J. H., Sutton, H. J., Cottrell, C. A., Phung, I., Ozorowski, G., Sewall, L. M., Nedellec, R., Nakao, C., Silva, M., Richey, S. T., Torre, J. L., Lee, W. H., Georgeson, E., Kubitz, M., Hodges, S., Mullen, T. M., Adachi, Y., Cirelli, K. M., Kaur, A., Allers, C., … Crotty, S. (2022). Long-primed germinal centres with enduring affinity maturation and clonal migration. Nature, 609(7929), 998–1004.

38. Heesters, B. A., Chatterjee, P., Kim, Y. A., Gonzalez, S. F., Kuligowski, M. P., Kirchhausen, T., & Carroll, M. C. (2013). Endocytosis and recycling of immune complexes by follicular dendritic cells enhances B cell antigen binding and activation. Immunity, 38(6), 1164–1175.

39. Kranich, J., & Krautler, N. J. (2016). How Follicular Dendritic Cells Shape the B-Cell Antigenome. Frontiers in immunology, 7, 225.

40. Hagemann, K., Riecken, K., Jung, J. M., Hildebrandt, H., Menzel, S., Bunders, M. J., Fehse, B., Koch-Nolte, F., Heinrich, F., Peine, S., Schulze Zur Wiesch, J., Brehm, T. T., Addo, M. M., Lütgehetmann, M., & Altfeld, M. (2022). Natural killer cell-mediated ADCC in SARS-CoV-2-infected individuals and vaccine recipients. European journal of immunology, 52(8), 1297–1307.

41. Beaudoin-Bussières, G., Chen, Y., Ullah, I., Prévost, J., Tolbert, W. D., Symmes, K., Ding, S., Benlarbi, M., Gong, S. Y., Tauzin, A., Gasser, R., Chatterjee, D., Vézina, D., Goyette, G., Richard, J., Zhou, F., Stamatatos, L., McGuire, A. T., Charest, H., Roger, M., … Finzi, A. (2022). A Fc-enhanced NTD-binding non-neutralizing antibody delays virus spread and synergizes with a nAb to protect mice from lethal SARS-CoV-2 infection. Cell reports, 38(7), 110368.

42. Bahnan, W., Wrighton, S., Sundwall, M., Bläckberg, A., Larsson, O., Höglund, U., Khakzad, H., Godzwon, M., Walle, M., Elder, E., Strand, A. S., Happonen, L., André, O., Ahnlide, J. K., Hellmark, T., Wendel-Hansen, V., Wallin, R. P., Malmstöm, J., Malmström, L., Ohlin, M., … Nordenfelt, P. (2022). Spike-Dependent Opsonization Indicates Both Dose-Dependent Inhibition of Phagocytosis and That Non-Neutralizing Antibodies Can Confer Protection to SARS-CoV-2. Frontiers in immunology, 12, 808932.

43. Masterman, K. A., Haigh, O. L., Tullett, K. M., Leal-Rojas, I. M., Walpole, C., Pearson, F. E., Cebon, J., Schmidt, C., O’Brien, L., Rosendahl, N., Daraj, G., Caminschi, I., Gschweng, E. H., Hollis, R. P., Kohn, D. B., Lahoud, M. H., & Radford, K. J. (2020). Human CLEC9A antibodies deliver NY-ESO-1 antigen to CD141+ dendritic cells to activate naïve and memory NY-ESO-1-specific CD8+ T cells. Journal for immunotherapy of cancer, 8(2), e000691.

44. Weiskopf, D., Angelo, M. A., de Azeredo, E. L., Sidney, J., Greenbaum, J. A., Fernando, A. N., Broadwater, A., Kolla, R. V., De Silva, A. D., de Silva, A. M., Mattia, K. A., Doranz, B. J., Grey, H. M., Shresta, S., Peters, B., & Sette, A. (2013). Comprehensive analysis of dengue virus-specific responses supports an HLA-linked protective role for CD8+ T cells. Proceedings of the National Academy of Sciences of the United States of America, 110(22), E2046–E2053.

45. Rivino, L., & Lim, M. Q. (2017). CD4+ and CD8+ T-cell immunity to Dengue - lessons for the study of Zika virus. Immunology, 150(2), 146–154.

46. Pérez-Gómez, A., Gasca-Capote, C., Vitallé, J., Ostos, F. J., Serna-Gallego, A., Trujillo-Rodríguez, M., Muñoz-Muela, E., Giráldez-Pérez, T., Praena-Segovia, J., Navarro-Amuedo, M. D., Paniagua-García, M., García-Gutiérrez, M., Aguilar-Guisado, M., Rivas-Jeremías, I., Jiménez-León, M. R., Bachiller, S., Fernández-Villar, A., Pérez-González, A., Gutiérrez-Valencia, A., Rafii-El-Idrissi Benhnia, M., … Virgen del Rocío Hospital COVID-19 and COHVID-GS Working Teams (2022). Deciphering the quality of SARS-CoV-2 specific T-cell response associated with disease severity, immune memory and heterologous response. Clinical and translational medicine, 12(4), e802.

47. Pennock, N. D., Gapin, L., & Kedl, R. M. (2014). IL-27 is required for shaping the magnitude, affinity distribution, and memory of T cells responding to subunit immunization. Proceedings of the National Academy of Sciences of the United States of America, 111(46), 16472–16477.

48. Gwyer Findlay, E., Villegas-Mendez, A., O’Regan, N., de Souza, J. B., Grady, L. M., Saris, C. J., Riley, E. M., & Couper, K. N. (2014). IL-27 receptor signaling regulates memory CD4+ T cell populations and suppresses rapid inflammatory responses during secondary malaria infection. Infection and immunity, 82(1), 10–20.

49. Kilgore, A. M., Pennock, N. D., & Kedl, R. M. (2020). cDC1 IL-27p28 Production Predicts Vaccine-Elicited CD8+ T Cell Memory and Protective Immunity. Journal of immunology (Baltimore, Md. : 1950), 204(3), 510–517.

50. Klarquist, J., Cross, E. W., Thompson, S. B., Willett, B., Aldridge, D. L., Caffrey-Carr, A. K., Xu, Z., Hunter, C. A., Getahun, A., & Kedl, R. M. (2021). B cells promote CD8 T cell primary and memory responses to subunit vaccines. Cell reports, 36(8), 109591.

51. Karsten, H., Cords, L., Westphal, T., Knapp, M., Brehm, T. T., Hermanussen, L., Omansen, T. F., Schmiedel, S., Woost, R., Ditt, V., Peine, S., Lütgehetmann, M., Huber, S., Ackermann, C., Wittner, M., Addo, M. M., Sette, A., Sidney, J., & Schulze Zur Wiesch, J. (2022). High-resolution analysis of individual spike peptide-specific CD4+ T-cell responses in vaccine recipients and COVID-19 patients. Clinical & translational immunology, 11(8), e1410.

52. Moss P. (2022). The T cell immune response against SARS-CoV-2. Nature immunology, 23(2), 186–193.

53. Lim, J. M. E., Tan, A. T., Le Bert, N., Hang, S. K., Low, J. G. H., & Bertoletti, A. (2022). SARS-CoV-2 breakthrough infection in vaccinees induces virus-specific nasal-resident CD8+ and CD4+ T cells of broad specificity. The Journal of experimental medicine, 219(10), e20220780.

54. Liu, J., Chandrashekar, A., Sellers, D., Barrett, J., Jacob-Dolan, C., Lifton, M., McMahan, K., Sciacca, M., VanWyk, H., Wu, C., Yu, J., Collier, A. Y., & Barouch, D. H. (2022). Vaccines elicit highly conserved cellular immunity to SARS-CoV-2 Omicron. Nature, 603(7901), 493–496.

55. Freeman, C. M., & Curtis, J. L. (2017). Lung Dendritic Cells: Shaping Immune Responses throughout Chronic Obstructive Pulmonary Disease Progression. American journal of respiratory cell and molecular biology, 56(2), 152–159.

56. Granot, T., Senda, T., Carpenter, D. J., Matsuoka, N., Weiner, J., Gordon, C. L., Miron, M., Kumar, B. V., Griesemer, A., Ho, S. H., Lerner, H., Thome, J. J. C., Connors, T., Reizis, B., & Farber, D. L. (2017). Dendritic Cells Display Subset and Tissue-Specific Maturation Dynamics over Human Life. Immunity, 46(3), 504–515.

57. Li, J., Ahmet, F., Sullivan, L. C., Brooks, A. G., Kent, S. J., De Rose, R., Salazar, A. M., Reis e Sousa, C., Shortman, K., Lahoud, M. H., Heath, W. R., & Caminschi I. (2015). Antibodies targeting Clec9A promote strong humoral immunity without adjuvant in mice and non-human primates. European journal of immunology, 45(3), 854–64.

57. Tullett, K. M., Leal Rojas, I. M., Minoda, Y., Tan, P. S., Zhang, J. G., Smith, C., Khanna, R., Shortman, K., Caminschi, I., Lahoud, M. H., & Radford, K. J. (2016). Targeting CLEC9A delivers antigen to human CD141^+^ DC for CD4^+^ and CD8^+^T cell recognition. Journal of clinical investigation insight, 1(7), e87102.

