## Supplemental Figures for "Single shot dendritic cell targeting SARS-CoV-2 vaccine candidate induces broad and durable systemic and mucosal immune responses"

### SUPPLEMENTARY FIGURES

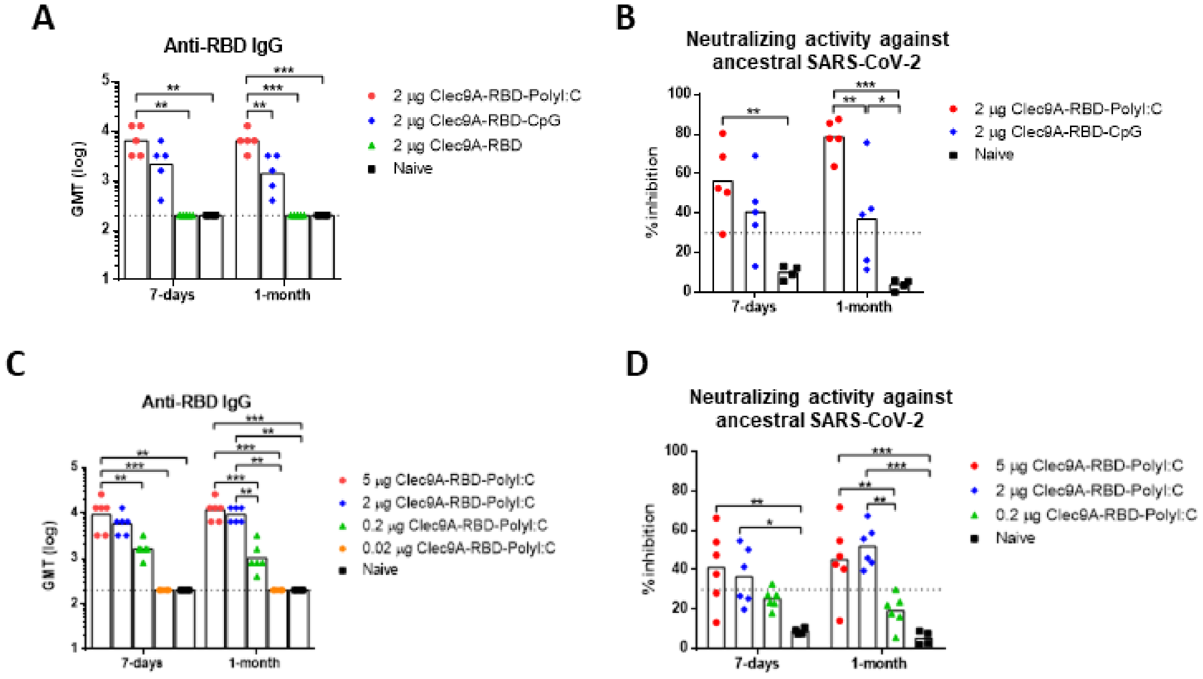


**Figure S1. Immunizing dose and adjuvant optimization.**

**(A, B)** 5-6 weeks old Balb/c mice were sc. immunized once with 2 µg Clec9A-RBD adjuvanted with 50 µg Poly I:C, 50 µg CpG, or no adjuvant. **(C, D)** 5-6 weeks old Balb/c mice were sc. immunized once with 5, 2, 0.2 and 0.02 µg of Clec9A-RBD adjuvanted with 50 µg Poly I:C. **(A, C)** SARS-CoV-2 RBD-specific IgG titers in immune sera were measured by ELISA. The dashed and dotted line represents the ELISA limit of detection (2.3 log). **(B, D)** Neutralizing activity of immune sera against ancestral SARS-CoV-2 was determined by c-Pass sVNT. The dashed line at 30% represents the cut-off value above which samples are considered positive. **(A-D)** Symbols represent individual animals and bars represent **(A, C)** geometric mean and **(B, D)** mean. Statistical analysis: **(A-D)** One-Way ANOVA with Tukey's correction for multiple comparisons. *p < 0.05, **p < 0.01, or ***p < 0.001.

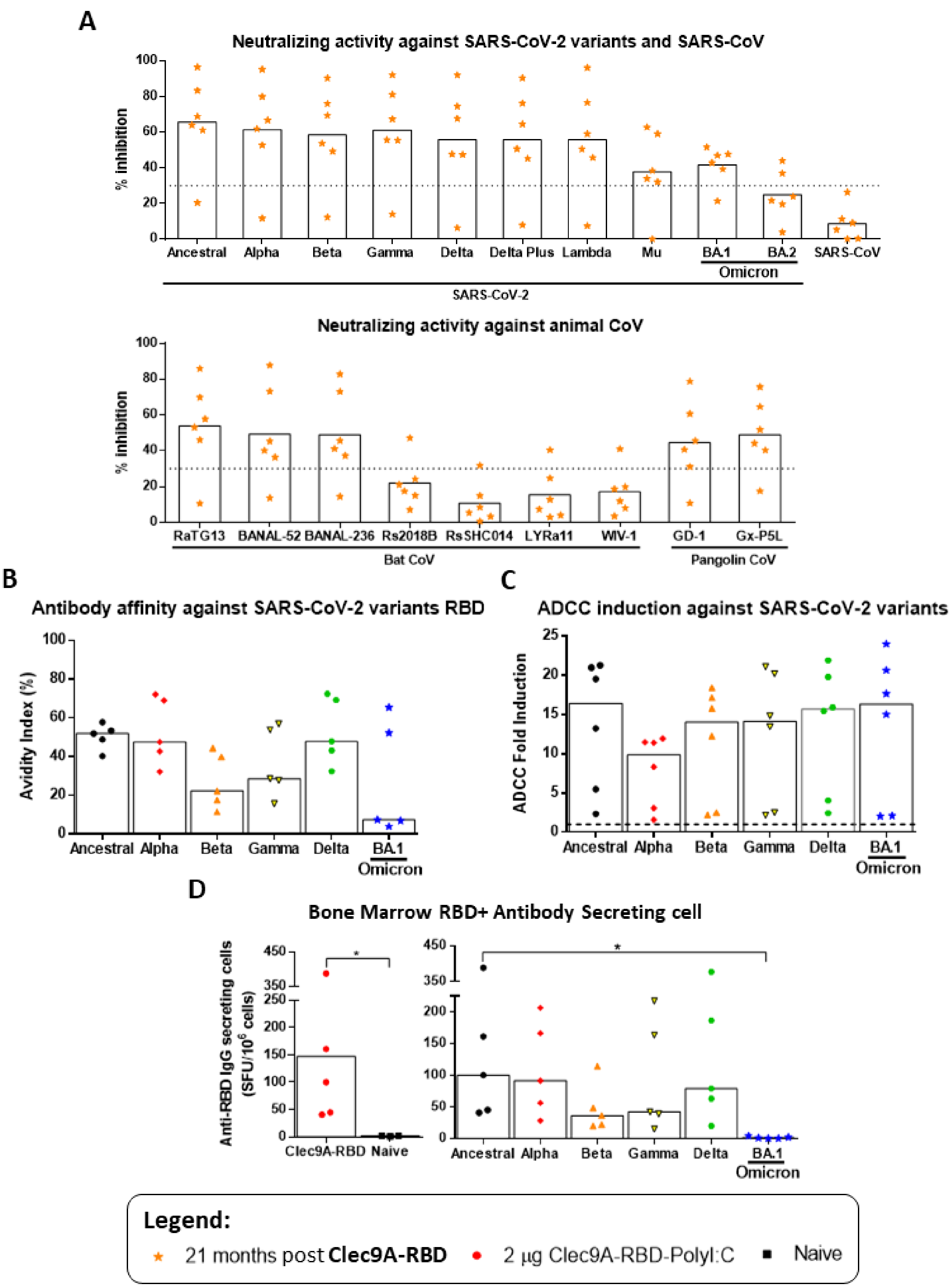


**Figure S2. RBD-specific humoral responses against ancestral SARS-CoV-2, SARS-CoV-2 variants and other sarbecoviruses at 21 months post-immunization with Clec9A-RBD.** 5-6 weeks old Balb/c mice were sc. immunized once with 2 µg Clec9A-RBD adjuvanted with 50 µg Poly I:C. Systemic humoral responses were analysed at 21 months post-immunization. **(A)** Serum neutralizing activity of the immune sera against a panel of 20 sarbecoviruses were determined via Multiplex sVNT. The dashed line at 30% represents the cut-off value above which samples are considered positive. **(B)** IgG avidity index and **(C)** ADCC activity of immune sera against ancestral and variant SARS-CoV-2 were determined by urea wash ELISA and ADCC Reporter Bioassay, respectively. The dashed line represents the baseline ADCC fold induction (1.0). **(D)** Frequency of bone marrow antibody secreting cells reactive to ancestral and variant SARS-CoV-2 RBD were enumerated by B cell ELISPOT. **(A-D)** Symbols represent individual animals and bars represent **(A)** mean and **(B-D)** median. Statistical analysis: **(D)** Mann Whitney test or Kruskal Walis test with Dunnett's correction for multiple comparisons. $*p < 0.05$.

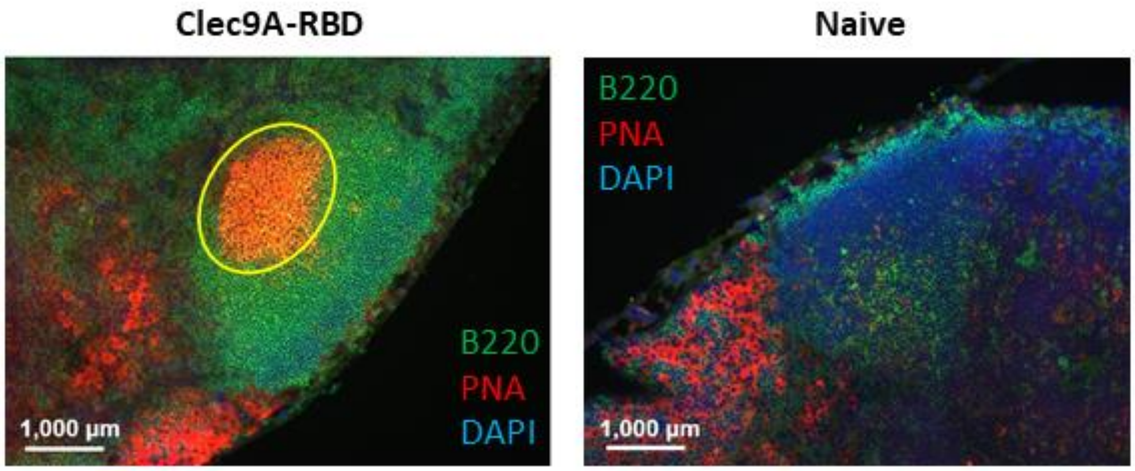


**Figure S3. Immune detection of GC B cells in brachial lymph nodes from Clec9A-RBD immunized mice.**

5-6 weeks old Balb/c mice were sc. immunized once with 2 µg Clec9A-RBD adjuvanted with 50 µg Poly I:C or left non immunized (naïve). Brachial lymph node sections harvested at 2 weeks post-immunization were stained with PNA, B220, and DAPI. GC B cells were defined as $B220^{+}$ $PNA^{+}$ (circled in yellow). Representative images are shown (scale bar: 1,000 µm).

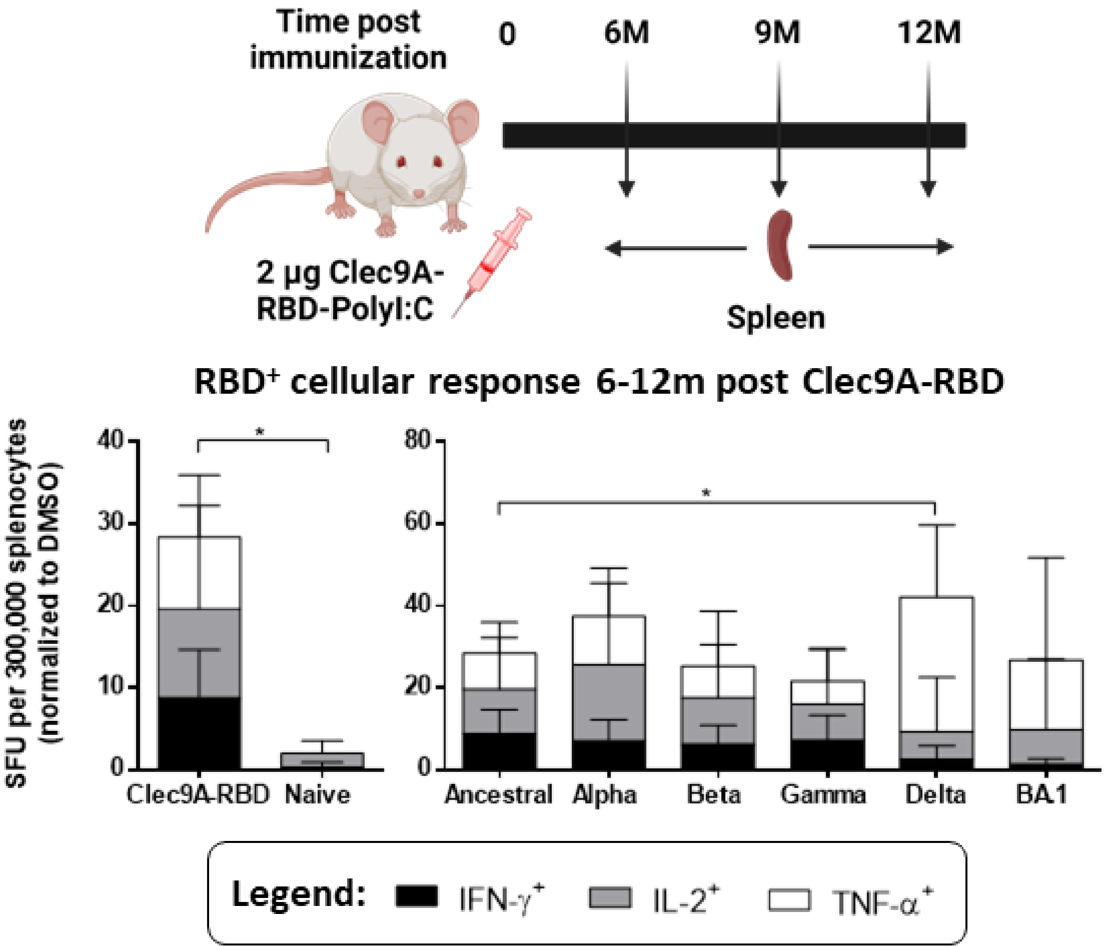


**Figure S4. RBD-specific cellular responses against ancestral and variant SARS-CoV-2 at 6-12 months upon single dose immunization with Clec9A-RBD.**

5-6 weeks old Balb/c mice (n=5) were sc. immunized once with 2 µg Clec9A-RBD adjuvanted with 50 µg Poly I:C, and spleens were harvested at 6-12 months post-immunization (n=2 at 6 and 9 months each, and n=1 at 12 months). The frequency of IFN-$\gamma^+$, IL-$2^+$, and TNF-$\alpha^+$ splenocytes was measured by IFN-γ/IL-2/TNF-α FluoroSPOT upon re-stimulation with ancestral and variant SARS-CoV-2 RBD peptides. Data were expressed as the mean ± SD. Statistical analysis: unpaired two tailed *t* test or One-Way ANOVA with Tukey's correction for multiple comparisons. *$p < 0.05$.

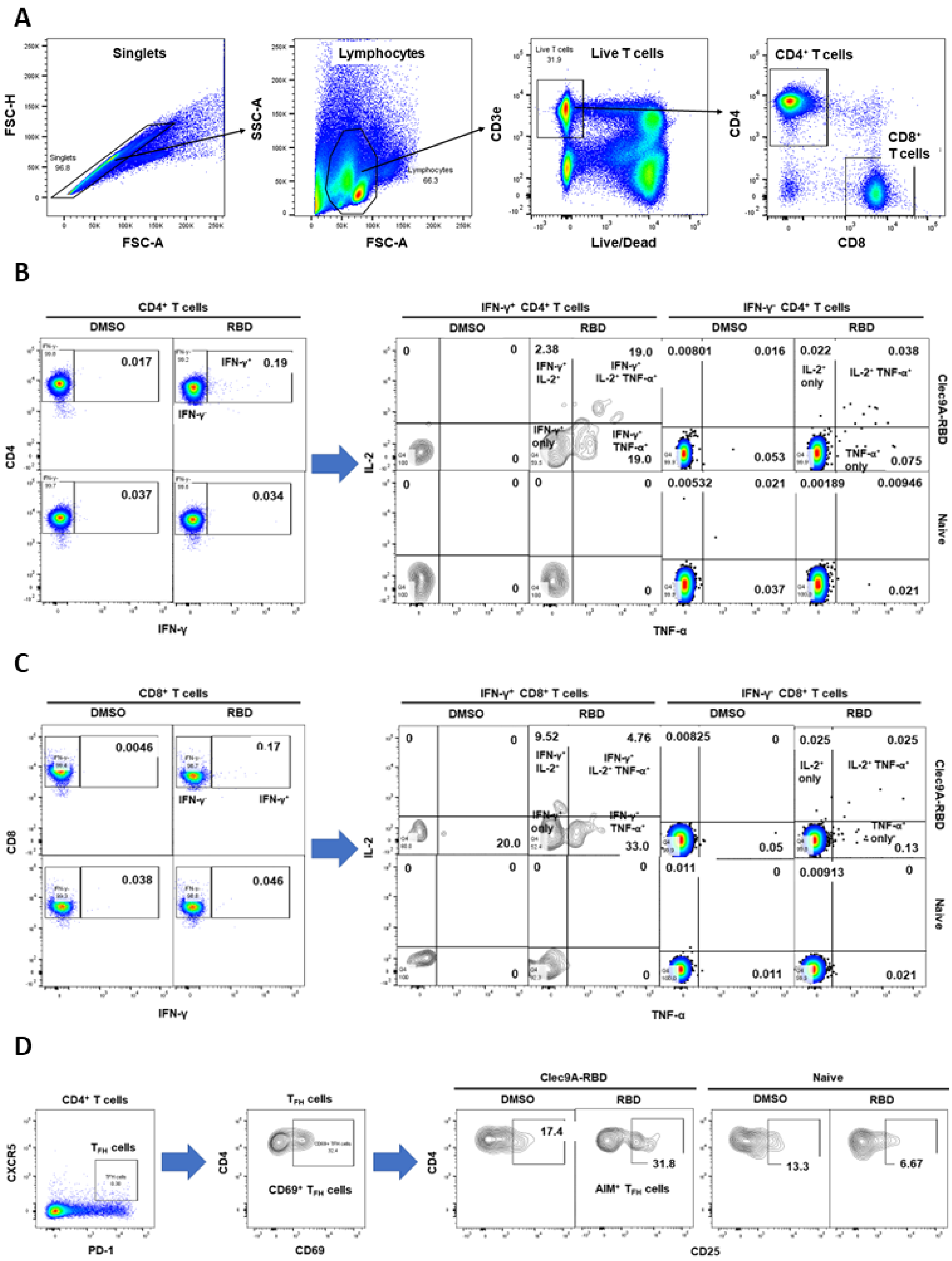


**Figure S5. Flow cytometry gating strategies to analyse $CD4^+$ and $CD8^+$ T cell subsets.**

5-6 weeks old Balb/c mice were sc. immunized once with 2 µg Clec9A-RBD adjuvanted with 50 µg PolyI:C. **(A)** Gating strategy of splenocytes from a representative Clec9A-RBD immunized and naïve mouse re-stimulated with ancestral SARS-CoV-2 RBD peptides to identify $CD4^+$ and $CD8^+$ T cell subsets. **(B)** Gating strategy to analyse $CD4^+$ T cell cytokine response. **(C)** Gating strategy to identify $CD8^+$ T cell cytokine response. **(D)** Gating strategy to identify $AIM^+$ $T_{FH}$ cells. Similar gating and analysis were performed for splenocytes re-stimulated with variant SARS-CoV-2 RBD peptides, and lung $CD4^+$ and $CD8^+$ T cells re-stimulated with ancestral SARS-CoV-2 RBD peptides.

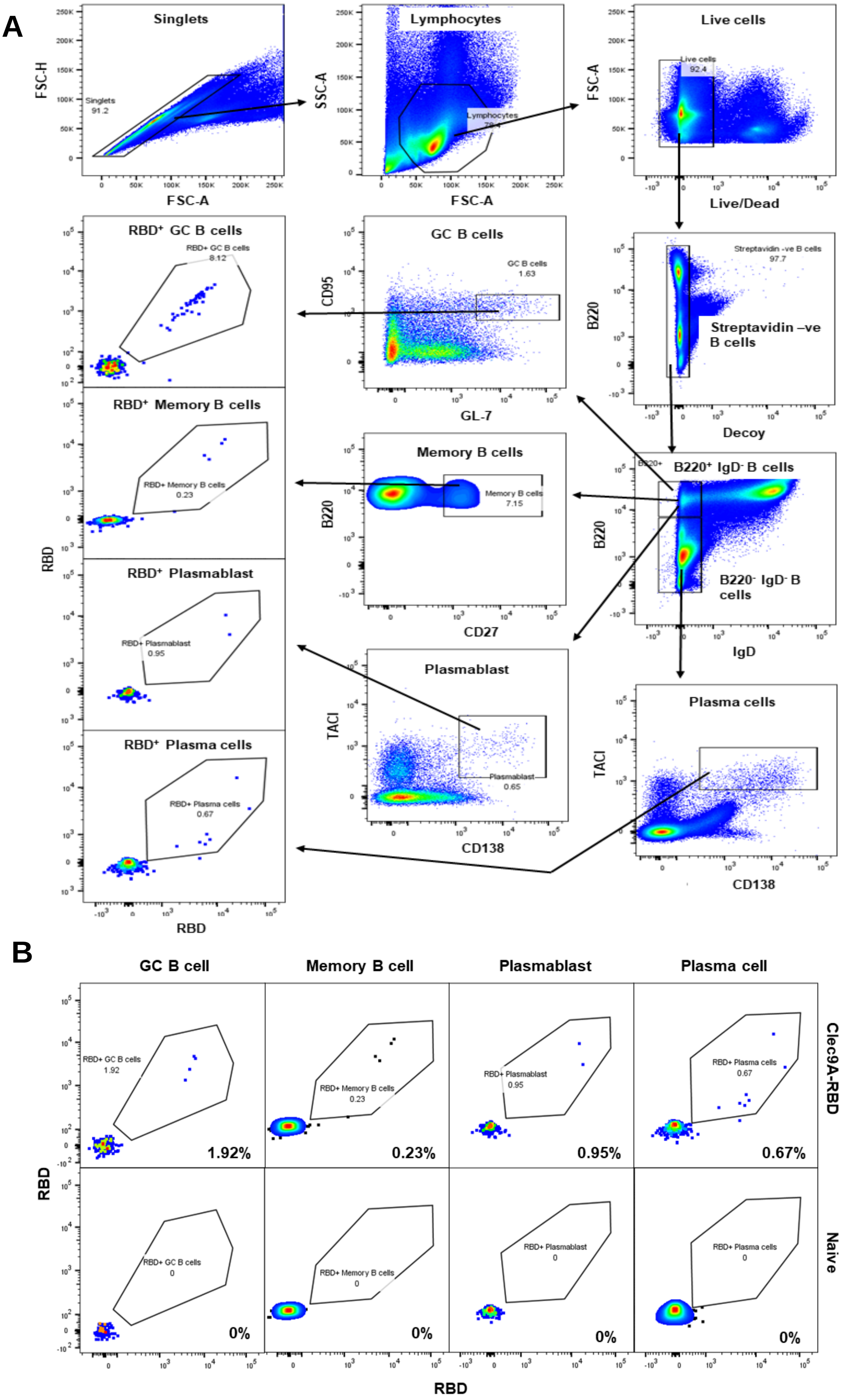


**Figure S6. Flow cytometry gating strategy to analyse B cell subsets.**

**(A)** Gating strategy to identify RBD-specific GC B cells, memory B cells, plasmablasts, and plasma cells. **(B)** Representative results of spleen RBD-specific GC B cells, memory B cells, plasmablasts, and plasma cells from Clec9A-RBD immunized and naïve mouse. Similar gating and analysis were performed to identify lung RBD-specific GC B cells.

**Table S1. RBD proteins from the 20 sarbecoviruses used in the Multiplex sVNT.**

| Clade | Sarbecovirus | Variant/Strain | Source |
|---|---|---|---|
| 1B | SARS-CoV-2 | Ancestral | Custom made by Genscript |
| 1B | SARS-CoV-2 | Alpha (B.1.1.7) | Custom made by Genscript |
| 1B | SARS-CoV-2 | Beta (B.1.351) | Custom made by Genscript |
| 1B | SARS-CoV-2 | Gamma (P.1) | Custom made by Genscript |
| 1B | SARS-CoV-2 | Delta (B.1.617.2) | Custom made by Genscript |
| 1B | SARS-CoV-2 | Delta Plus (B.1.617.2.1) | Produced in-house in HEK293T cells as described previously [23] |
| 1B | SARS-CoV-2 | Lambda (C.37) | Produced in-house in HEK293T cells as described previously [23] |
| 1B | SARS-CoV-2 | Mu (B.1.621) | Produced in-house in HEK293T cells as described previously [23] |
| 1B | SARS-CoV-2 | BA.1 (Omicron) | SPD-C522k; Acrobiosystems, Newark, Delaware, USA |
| 1B | SARS-CoV-2 | BA.2 (Omicron) | Produced in-house in HEK293T cells as described previously [23] |
| 1B | Bat CoV | RaTG13 | Custom made by Genscript |
| 1B | Bat CoV | BANAL-52 | Produced in-house in HEK293T cells as described previously [23] |
| 1B | Bat CoV | BANAL-236 | Produced in-house in HEK293T cells as described previously [23] |
| 1B | Pangolin CoV | GD-1 | Produced in-house in HEK293T cells as described previously [23] |
| 1B | Pangolin CoV | GX-P5L | Custom made by Genscript |
| 1A | SARS-CoV | SARS-CoV | Custom made by Genscript |
| 1A | Bat CoV | Rs2018B | Produced in-house in HEK293T cells as described previously [23] |
| 1A | Bat CoV | RsSHC014 | Produced in-house in HEK293T cells as described previously [23] |
| 1A | Bat CoV | LYRa11 | Produced in-house in HEK293T cells as described previously [23] |
| 1A | Bat CoV | WIV-1 | Produced in-house in HEK293T cells as described previously [23] |

**Table S2. Antibodies used for flow cytometry staining of intracellular cytokines, AIM$^+$ T$_{FH}$ cells, and B cell subsets.**

| Surface/Intracellular | Marker | Fluorophore | Dilution | Product Number |
|---|---|---|---|---|
| **Intracellular cytokine staining** | | | | |
| Surface | CD3 | Alexa Fluor 488 | 1:200 | 100210; Biolegend |
| | CD4 | BUV395 | | 563790; BD Biosciences |
| | CD8 | BV711 | | 563046; BD Biosciences |
| Intracellular | IFN-γ | BV421 | | 505829; Biolegend |
| | IL-2 | PE | | 503807; Biolegend |
| | TNF-α | APC | | 506307; Biolegend |
| **AIM$^+$ T$_{FH}$ cells** | | | | |
| Surface | CD3 | Alexa Fluor 488 | 1:200 | 100210; Biolegend |
| | CD4 | BUV395 | | 563790; BD Biosciences |
| | CXCR5 | APC | | 145506; Biolegend |
| | PD-1 | PE | | 568261; BD Biosciences |
| | CD25 | BV510 | | 563037; BD Biosciences |
| | CD69 | BV421 | | 562920; BD Biosciences |
| **B cell subsets** | | | | |
| Surface (pan B cell) | B220 | BUV395 | 1:200 | 563793; BD Biosciences |
| | IgD | BV711 | | 564275; BD Biosciences |
| Surface (GC B cell) | GL-7 | Alexa Fluor 647 | | 561529; BD Biosciences |
| | CD95 | BV605 | | 152612; Biolegend |
| Surface (Memory B cell) | CD27 | | | 563365; BD Biosciences |
| Surface (Plasmablast/Plasma cell) | CD138 | | | 563147; BD Biosciences |
| | TACI | Alexa Fluor 647 | | 558453; BD Biosciences |
